# Gene expression noise is reduced in communicating synthetic cell populations

**DOI:** 10.64898/2026.09.24.754039

**Authors:** Naresh Yadrapalli, David T. Gonzales, T-Y Dora Tang

## Abstract

A major goal in bottom-up synthetic biology is the construction of multicellular synthetic systems capable of coordinated and robust collective behaviours. However, robustness is often limited by noise and variability arising from increased molecular complexity. Whilst communication has been implemented in synthetic multi-cellular systems, the ability for communication to suppress cell free gene expression variability in populations of synthetic cells remain unexplored. To address this, we encapsulated the Lux and Las quorum sensing gene circuits in lipid vesicles under cell-free conditions to test the effect of communication on reducing cell-free gene expression variability across the population. Our results show that communication, limiting expression resources, and membrane surface effects can reduce gene expression variability. Resource limited Gillespie simulations for transcription and translation show that communication-mediated coupling reduces population-level expression noise under constrained and excess resource conditions. Together, our work provides simple strategies to reduce gene expression variability and thereby improve robustness in synthetic multicellular systems, an important criteria for the future applications of synthetic cells.

## Introduction

One of the major goals of bottom-up synthetic biology is the construction of multicellular synthetic systems capable of coordinated and robust collective behaviours^1–3^. Lipid vesicles that contain DNA and transcription-translation machinery have been established as the primary building block for synthetic cellular systems and could be considered as a rudimentary synthetic cell^4^. These platforms have been engineered for communication^5^; sensing^6^; control of gene expression^7^; DNA replication^8^ coupled to cell division^9^; drug delivery^10^. However, a major limitation of cell-free lipid vesicles is gene expression noise that could compromise the robustness of synthetic systems^11,12^. Here, gene expression noise is defined as cell-to-cell variability within a genetically identical population^13,14^. In biological systems, stochastic gene expression contributes to phenotypic heterogeneity, influencing processes such as cell fate determination, stress adaptation, antimicrobial resistance and cancer progression^14–16^. This variability could arise from heterogeneous distributions of molecular content across the cells and/or from the random nature of chemical reactions^16^. There are a number of strategies which can be used to minimize variability in synthetic cellular systems. Microfluidic approaches can reduce variability by tightly regulating the distribution of chemical content across a population of vesicles^7,17^. Additional strategies to control gene expression noise, that have been developed in cells include modifications to the promoter site ^18^; tuning the gene circuit design^19^; integration of regulatory features^20^. In addition, communication provides an alternative strategy to regulate cells across a population. In living systems, communication enables cells to co-ordinate responses across populations, reduce stochastic fluctuations and perform complex functions ranging from microbial quorum sensing to tissue development^21,22^. Furthermore, quorum sensing bacteria co-ordinate gene expression through diffusible signaling molecules^22^. However, the ability for communication to suppress cell-free gene expression variability in population of vesicles remains largely unexplored. To this end, we test whether communication can provide a rational strategy for reducing gene expression noise within populations of rudimentary synthetic cells.

Communication has been well established in synthetic ^23,24^ with bacterial ^25^ systems where autocrine, paracrine, juxtacrine communication has been demonstrated^2^. To achieve communication, researchers implemented different strategies including the integration of cell-free gene expression^23^; quorum-sensing circuits^25,26^; enzymatic signal amplification^27^; membrane pores for controlled molecular exchange^23^; and DNA/RNA-based signaling^28^. More recent work has extended these platforms to bidirectional signaling^24^, distributed computation and feedback across synthetic-cell populations^28^ and spatially controlled synthetic tissues^29,30^, demonstrating increasingly complex collective behaviors. However, these studies have largely focused on establishing communication and signal-processing function and have not considered the role of communication to suppress cell-to-cell variability in cell-free gene expression across synthetic-cell population. It is important to control the variability in cellular functions to achieve collective synchronous responses within synthetic cells.

To this end, we implemented an on-chip inverted emulsion platform^31^ to facilitate the concurrent generation of synthetic cells with communication and imaging. In this instance synthetic cells are lipid vesicles containing transcription-translation machinery. To achieve communication, we encapsulated Lux and Las quorum sensing gene circuits into lipid vesicles under cell-free conditions to induce autocrine and paracrine communication, for three (Lux system) and six plasmid gene circuits (Lux and Las system)^25^. The measured fluorescence intensity of the expressed reporter protein was used to determine the variability of gene expression across the population. Our results show that gene expression variability can be reduced by communication; limiting the resources and by tuning the composition of the membrane. We further confirmed these results using a Gillispie stochastic simulation algorithm^32^. Our work provides simple strategies to reduce gene expression variability and thus increase robustness in synthetic multi-cellular systems.

## Results

To confirm that the gene circuits were compatible with lipid vesicles, we assembled lipid vesicles ranging from 5-40 microns by the inverted emulsion method^31^ (Supplementary figure 1). The lipid vesicles contained home-made bacterial extracts that drive cell-free transcription–translation with its corresponding resources and energy (see Materials and methods in SI) and a 6 plasmid-2 node quorum sensing gene circuit^24^ (Supplementary figure 1). The gene circuit couples the LuxI/LuxR and

LasI/LasR modules such that LuxI is constitutively expressed to synthesise AHL N-(3-oxohexanoyl) homoserine lactone (3OC6-HSL). This will activate plasmids coding for a reporter protein eGFP and a LasI via binding with constitutively expressed Lux R and promoter activation. Las I drive the production of a second signaling molecule AHL N-(3-oxododecanoyl)homoserine lactone (3OC12-HSL)) that activates constitutively expressed LasR to express mScarlett-1 as previously described (Supplementary figure 1)^24^. Confocal imaging confirmed that the lipid vesicles were stable and expressed eGFP and mScarlet-I for Lux I/LuxR module and the LasI/LasR module respectively (Supplementary figure 1, Movie S1).

Having confirmed that the cell-free gene circuit was compatible with lipid vesicles, we compared the effect of autocrine and paracrine communication on population variability to obtain CV^2^ and study the effect of communication on population variability. The three plasmid Lux module (pT7-LuxI, pT7-Lux R and pLux-eGFP) were encapsulated into one vesicle or split across two vesicles (Figure 1). In the latter case, the first population contained pT7-Lux I plasmid and the second vesicle population contained pT7-Lux R and pLux-eGFP (Figure 1A). In the single vesicle case, the 3-plasmid gene circuit will allow both autocrine and juxtacrine (adjacent cells) communication, we term this system autocrine (nc-3). In the case of the two populations, reporter protein expression is induced by paracrine communication via 3OC6-HSL. This system is termed paracrine (c-3). To compare the effect of communication, each plasmid was encapsulated at a concentration of 4 nM. For paracrine communication, a two-chambered chip design was used to produce two populations of vesicles with the required plasmids, connected via a 17 mm long channel (Supplementary figure 2). This specific chip design principle allows for simultaneous production, communication and visualisation (due to the glass bottom) of the produced vesicles without any perturbation. Confocal imaging of the vesicles showed positive expression of eGFP in both the communicating and non-communication case (Figure 1B, Supplementary figure 3, Movie S2). Time lapse imaging showed, for the case of autocrine (nc-3), eGFP expression reached steady state after 6 hours whilst the steady state was attained after 4 hours in the case of the paracrine (c-3) system (Supplementary figure 4). Comparisons of the fluorescence intensity range of each vesicle across the populations (Figure 1Ci) and its respective CV^2^ (Figure 1Cii) (see also methods), at steady state, revealed that communication led to a decrease in the gene expression variability across the population by more than three times. In the case of cell-free expression in lipid vesicles, it has been well documented that gene expression variability across the population could arise from variable encapsulation of molecular components where the distribution of proteins, plasmid DNA and resources are not equally distributed across the population^33^. To determine whether the noise from encapsulation is a major factor for gene circuit expression variability we undertook control experiments that measure the variation in the encapsulation efficiency of purified eGFP protein (100 nM) and of eGFP expression alone. For the latter case, we encapsulated 4 nM of pT7 eGFP into lipid vesicles (Figure 1D and Supplementary figure 5). Analysis of the gene expression variability across the vesicle population showed low levels of noise: for purified protein CV^2^ ~0.06 and for eGFP expression-CV^2^ ~0.1. In this case CV^2^ was lower in our control experiments compared to populations containing the gene circuit. Here, autocrine (nc-3) exhibited greater variability (CV^2^ ~0.3) compared to expression of a single plasmid (CV^2^ ~0.1). In the case of paracrine (c-3), the noise is reduced (CV^2^ ~0.05) compared to autocrine (nc-3) but is comparable to the control experiments. Taken together, this indicates that the addition of the Lux system to the reaction mix increases the noise compared to a single plasmid while communication is reducing population variability.

**Figure 1:**
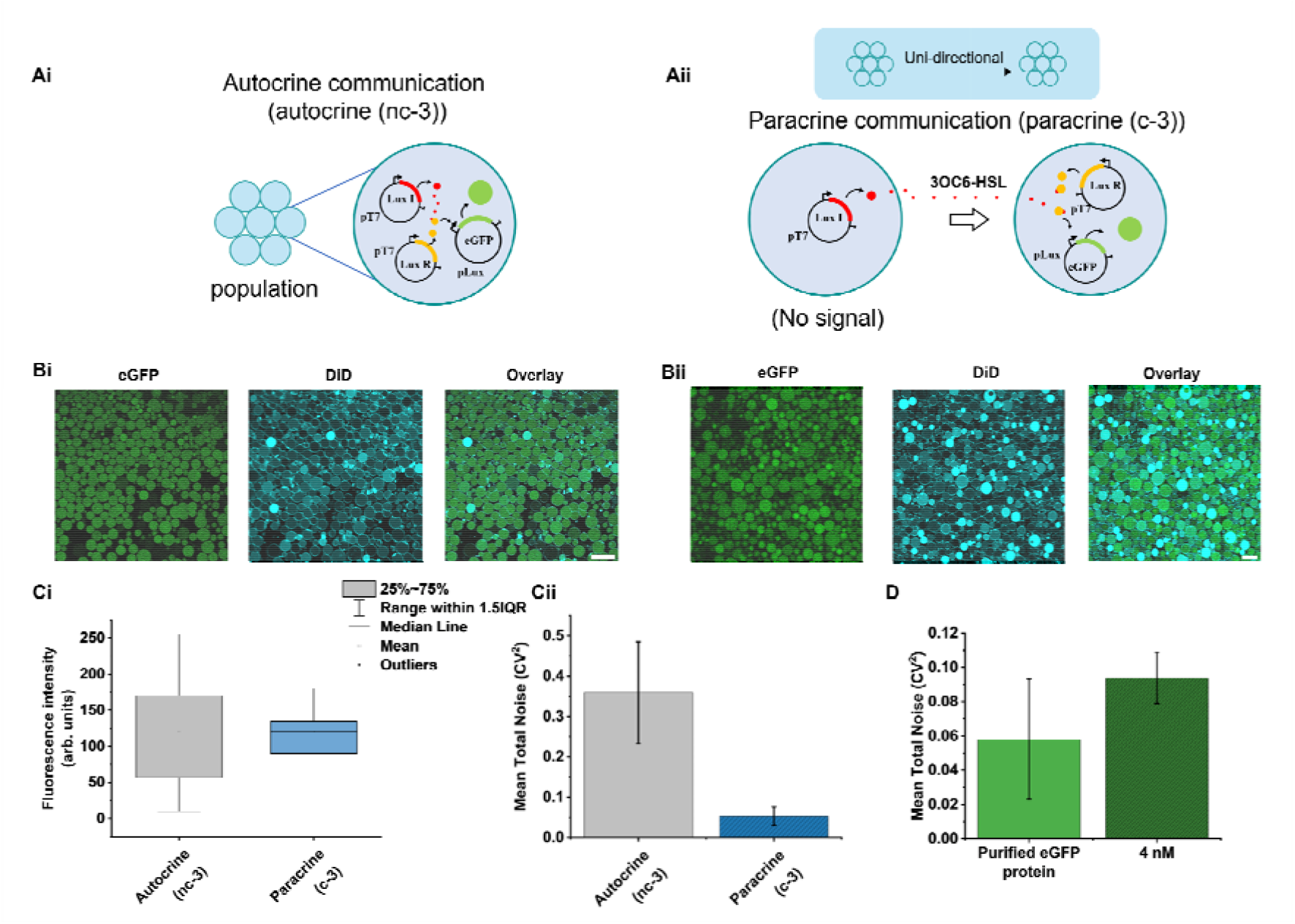
Paracrine communication reduces gene-expression variability in synthetic-cell populations. **(Ai)** Schematic showing the autocrine three-component circuit, in which LuxI, LuxR, and pLux-eGFP are encapsulated within the same synthetic cells (autocrine nc-3). Locally produced 3OC6-HSL drives eGFP expression from the pLux promoter. **(Aii)** Schematic of the unidirectional paracrine communication system (paracrine c-3), in which sender synthetic cells produce 3OC6-HSL but do not contain reporter plasmid, while receiver synthetic cells contain LuxR and pLux-eGFP, thus respond to the diffusible signal. **(Bi–Bii)** Representative confocal images showing eGFP reporter expression, DiD-labelled membranes, and merged channels showing expression contained within the lumen of the synthetic cells for both autocrine and paracrine populations. **(Ci)** Distribution of endpoint eGFP fluorescence intensities in autocrine and paracrine synthetic cells. Boxes indicate the 25–75% interquartile range, horizontal lines mark the median, open squares the mean, whiskers extend to 1.5× IQR, and symbols denote outliers. **(Cii)** Mean total expression noise, quantified as CV^2^, showing reduced population-level variability in the paracrine receiver system compared with the autocrine circuit. **(D)** Control comparison of total fluorescence noise for encapsulated purified eGFP protein and 4 nM pT7 eGFP expression condition. Error bars represent variation across replicate measurements, n = 4. Scale bars, 50 µm.

To confirm that communication alone would reduce gene expression variability, we compared CV^2^ for non-communicating and communicating populations of vesicles containing the Lux-Las six plasmid gene circuit. For the autocrine non-communicating population (autocrine (nc-6)), we analysed vesicles that exhibited fluorescence (Figure 2Ai). For the paracrine communicating system (paracrine (c-6)), the six plasmids were split to enable bi-directional communication with the pT7-LuxI, pT7 LasR and pLas-eGFP in vesicles within population 1 and pT7-LuxR, pLux-eGFP and pLux LasI in vesicles in population 2 that allowed the reporting of Lux and Las modules via two-way communication along the channel between the populations (Figure 2Aii, Supplementary figure 6). Again, confocal imaging showed fluorescence intensity in the lipid vesicles from eGFP expression reached steady state after 6 hours of expression (Figure 2B and Supplementary figure 7). Analysis of the individual vesicles across the population showed that the range in fluorescence intensities (Figure 2Ci) and the noise (Figure 2Cii) were reduced by approximately 5 times by communication. Our result confirms that communication provides a handle to reduce population variability and increase the robustness of population expression output.

**Figure 2:**
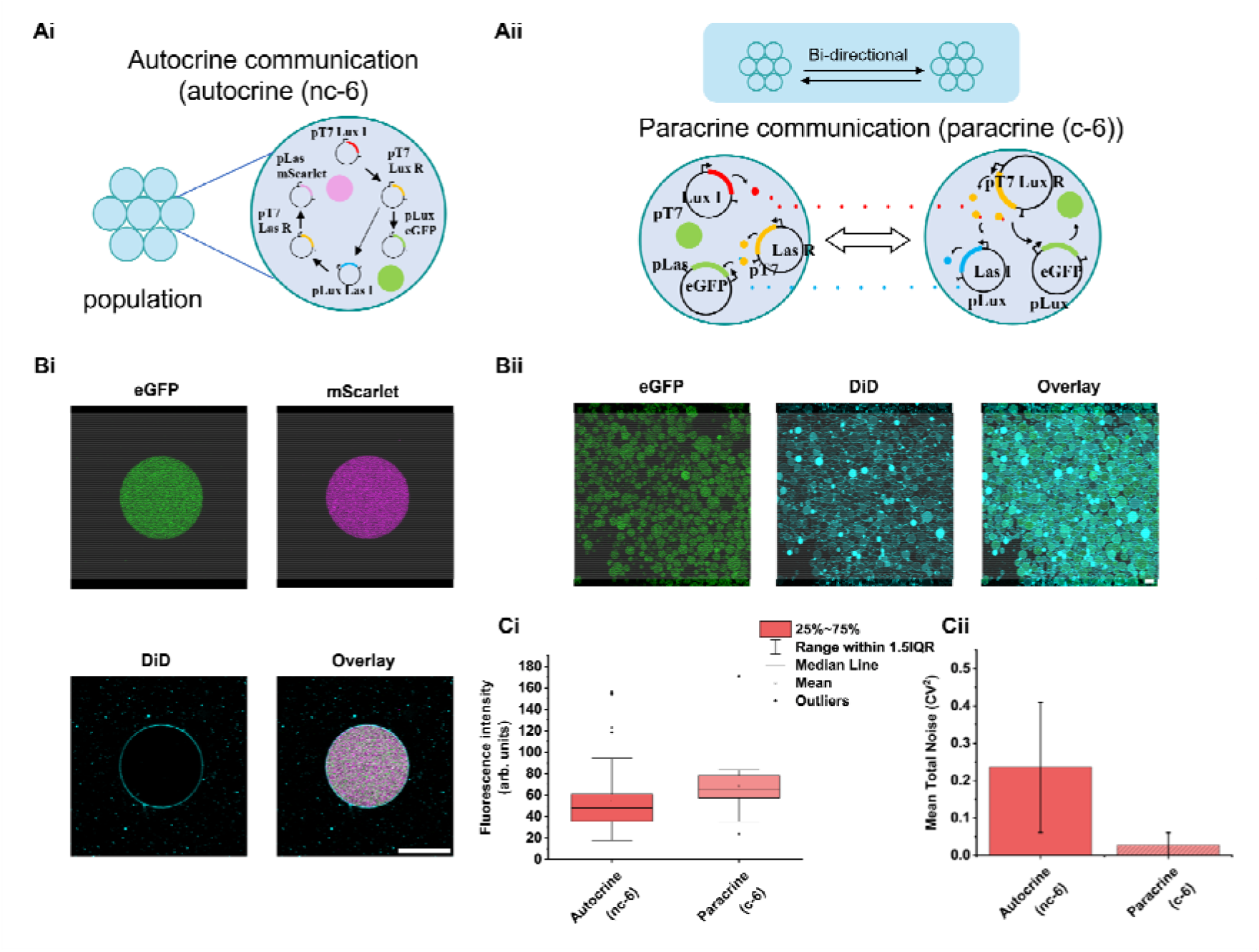
Bidirectional paracrine communication reduces expression variability in six-component synthetic-cell circuits. **(Ai)** Schematic of the autocrine six-component circuit (autocrine (nc-6)), in which LuxI/LuxR and LasI/LasR quorum-sensing modules are encapsulated within the same synthetic cells to drive pLux-eGFP and pLas-mScarlet-I expression. **(Aii)** Schematic of the bidirectional paracrine communication architecture (paracrine (c-6)), in which two synthetic cell populations exchange diffusible Lux and Las AHL signals to activate reporter expression in neighbouring receiver populations. **(Bi)** Representative confocal images of an autocrine synthetic cell showing eGFP, mScarlet-I, DiD-labelled membrane, and merged channels with expression colocalized within the vesicle. Scale bar, 20 µm. **(Bii)** Representative confocal images of the paracrine synthetic-cell population showing eGFP expression, DiD-labelled membranes, and overlay showing the expression within the confined region of the vesicles. Scale bar, 20 µm. **(Ci)** Endpoint reporter fluorescence distributions for autocrine and paracrine six-component circuits. Boxes indicate the 25–75% interquartile range, horizontal lines mark the median, open squares the mean, whiskers extend to 1.5× IQR, and symbols denote outliers. **(Cii)** Mean total expression noise, quantified as CV^2^, showing reduced population-level variability in the bidirectional paracrine architecture compared with the autocrine circuit. Error bars represent variation across replicate measurements, n = 4.

Whilst it can be challenging to provide a definitive explanation for communication driven noise reduction in gene expression, one could consider the effect of plasmid concentration as an important factor. In our experiments, we note that the concentration of plasmids in the vesicles will be greater in the case of autocrine (nc-3) (3 plasmid types) case compared to reporter protein expression population in paracrine (c-3) (2 plasmid types). In the former case, the individual plasmids are incorporated at 4 nM which means that a single vesicle will contain 12 nM of plasmid. For the latter, the vesicles containing pT7-LuxI will have a final concentration of 4 nM and the vesicles in the second population (reporter expressing) will contain 8 nM of plasmid. This may have a significant effect on the production and concentration of the lactone. Unlike the autocrine (nc-3) condition, in the paracrine (c-3) condition, there are no competing reactions for production of the signaling molecule (LuxI) within vesicles of population one, while the resources are identical across both systems. Thus, the availability of the resources is higher, thus the amount of signaling molecules produced. Similarly, within the vesicles of population two, for the production of eGFP (total plasmid concentration 8 nM) will be greater than the autocrine (nc-3) alternative (total plasmid concentration 12 nM). This could lead to a more rapid production of both the signaling molecule and thus reporter protein in the paracrine (c-3) case. Indeed, fitting the population averaged trajectories with a four-parameter Boltzmann (logistic) function with adjusted R^2^ values of 0.996 provided the ability to make qualitative comparisons between the experiments (Supplementary figure 8). For the autocrine (nc-3) experiments the time to half-maximum intensity of eGFP was achieved after 2.3 h with a steep rise characterised by a 0.45 h time constant. In comparison, for paracrine (c-3), the pLux-GFP response in receiver vesicles was markedly faster, achieving half-maximal activation within 1.23 h with a broader rise time (~0.7 h). This is consistent with faster 3OC6-HSL production and response to the lactone signal that can be attributed to the absence of the competition of resources from the pT7-LuxI.

It was interesting to note that when comparing the noise between the three plasmid and six plasmid gene circuits for the autocrine (nc) case there was a 1.5 fold decrease in CV^2^ (Supplementary figure 9). In the case of paracrine communication, we observed a 2.5 fold decrease from the three plasmid gene circuit to the six plasmid gene circuit (Supplementary figure 9). This suggests that as competition for resources can lead to a reduction of noise. To test this, we systematically changed the plasmid concentration from 0.5 nM to 4 nM of each plasmid for three and six plasmid gene circuit that undergoes paracrine communication (Figure 3B, Supplementary figure 10). We opted to test the paracrine over the autocrine communicating system as this system had the overall lower noise. Our results showed a decrease in the variability with increasing plasmid concentration for both paracrine (c-3) and paracrine (c-6) cases. For paracrine (c-3), the total noise decreased almost 5-fold with an 8 times difference in plasmid concentration for LuxI/LuxR–pLux–eGFP (Figure 3AI, Supplementary figure 11). This dropped from ~0.5 CV^2^ at 0.5 nM DNA to <0.1 CV^2^ at 4 nM. A similar trend is apparent in the six plasmid gene circuit (Figure 3Aii, Supplementary figure 12) where ~0.5 CV^2^ at 0.5 nM DNA to <0.1 CV^2^ at 4 nM. In addition, pLas generally exhibits a slightly higher CV^2^ than pLux at the same DNA dosage (Figure 3Aii). This could be attributed to the differences in variability from the pLas and pLux promoter, non-linear competition for resources between the plasmids or from variability that arises down-stream of the promoter that are different for the Las and Lux modules. Nevertheless, our results clearly show that population expression variability is dependent on the balance between transcription, translation and the amount of resources available.

**Figure 3.**
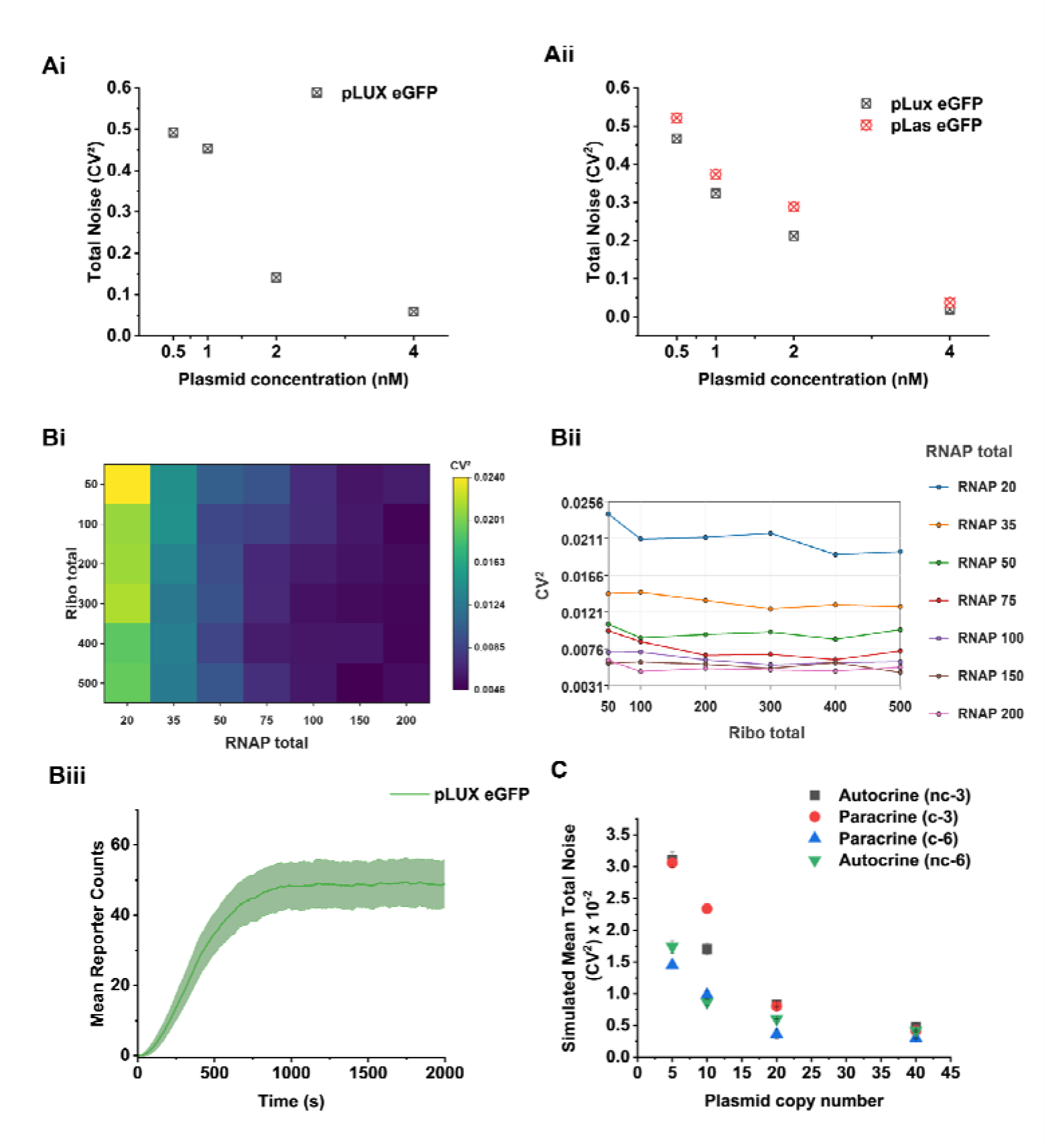
Plasmid and resource copy dependent suppression of gene-expression noise. **(Ai)** Experimental evidence of change in gene expression noise (CV^2^) for pLux-eGFP as a function of plasmid concentration in the three-component paracrine circuit. Increasing plasmid concentration progressively reduced reporter noise. **(Aii)** Experimental evidence for pLux-eGFP and pLas-eGFP in the six-component paracrine circuit, showing reduced noise at higher plasmid concentrations for both reporter outputs. n = 4 **(Bi)** Heat map showing simulated (CV^2^) across different total RNAP and ribosome abundances, indicating higher noise under low-resource conditions and reduced noise as expression resources increase. **(Bii)** Line profiles of (CV^2^) as a function of ribosome abundance at different RNAP levels, showing that RNAP availability strongly influences expression variability. **(Biii)** Representative simulated kinetic trace of mean pLux-eGFP reporter expression over time; shaded region indicates variability around the mean (N, 500). **(C)** Comparison of simulated mean total noise across autocrine and paracrine circuit architectures as a function of plasmid copy number. Across gene circuits, increasing plasmid copy number reduced CV^2^, supporting the experimental observation that communication can suppress gene-expression variability in synthetic-cell populations.

To further substantiate our results that limiting the resources reduces the variability, we performed Gillespie stochastic simulations based on mass action kinetics that include finite resource constraints and inter-compartment diffusion as described by Goetz et. al. 2025^32^ (Figure 3B, Supplementary figure 13). The kinetic profiles were simulated for the average GFP concentration, standard deviations and CV^2^. Simulations were performed for the four different experimental setups (Figure 1A and 2A). For the simulation, it was important to define a plasmid copy number so that all systems operate within their dynamic range and pLux–eGFP achieved similar mean amplitudes. When the total noise is plotted for this common condition (Figure 3B), gene circuits that distribute the resource pool over different plasmid types and greater plasmid copy numbers show systematically lower noise in pLux–eGFP. In general, we observe the same qualitative general trends as those observed from our experiments. Increasing the plasmid copy number leads to a decrease in the simulated noise. At low plasmid copy number (5), there is a greater difference in variability across the experiments where the three plasmid gene circuit is more noisy than the six plasmid gene circuit (Figure 3C). In the case of a plasmid copy number of ten, simulations show a lower noise for autocrine communication compared to paracrine. The overall observation is that, under resource limited condition, increasing plasmid copy number reduces the noise. While it is challenging to achieve resource variation without the effects of dilution in experimental setting, simulations are devoid of such constraints. To examine how specific resources such as ribosomes (ribo) and RNAP dependence at a fixed plasmid copy number (10) alter noise, more broadly, we next swept RNAP/ribosome resource availability simulations. The resource sweep further indicated that transcriptional resource availability was the dominant determinant of simulated noise. Across all four circuit architectures, increasing RNAP abundance (20 to 200) produced a much larger reduction in CV^2^ (Figure 3B and C and Supplementary figure 13) than increasing ribosome abundance (50 to 500), suggesting that transcriptional fluctuations and promoter-level competition are the primary contributors to variability. Ribosome availability had a weaker but still detectable effect, particularly in the more resource-demanding six-plasmid architecture. Thus, within the simulated parameter regime, RNAP limitation rather than ribosome limitation appears to be the main resource bottleneck controlling noise. Taken together, the simulations qualitatively agree with our experimental results that limited resources play a role in reducing population noise.

To further identify parameters that can contribute to noise reduction, we compared the CV^2^ between all four conditions (autocrine and paracrine systems with 3 and 6 plasmids). Previous research had shown an inverse scaling law between noise and size, ^34,35^ thus we looked at the effect of size against gene expression variability across our population architectures. Plotting the fluorescence output against the size of the vesicle shows a negative correlation (Supplementary figure 14). Thus, correcting for the size of the vesicle, the variability is reduced by at least 10 % (see methods and Figure 4A). All the four systems can also be categorized under equi-resource and equi-plasmid concentration to further understand the role of resources-DNA-size on noise (Supplementary figure 15). When comparing the equi-resource 3 plasmid-type condition, between the autocrine and paracrine systems, a narrower distribution of endpoint eGFP intensities is observed in the paracrine system compared to the non-communicating / autocrine systems. A correspondingly lower mean total noise, both before and after correcting for vesicle-size effects is evident (Figure 4A). Furthermore, with vesicle populations with an equi-resource and equi-plasmid concentrations (i.e., pLux GFP expressing autocrine nc-3 and paracrine (c-6)) showed reduction in noise levels after size adjustment. The size adjustment for paracrine communication, shows reduction in the variability by 58% (Figure 4B). This suggests that surface effects can play a role in reducing variability. To test this, we included 1 mol % of pegylated lipids for the preparation of the lipid vesicles that contain all the six plasmid gene circuit with pLux-GFP and pLas-Scarlet (Figure 4C). The addition of pegylated lipids shows a 40 % reduction in the variability. This shows that reducing surface effects between the membrane and the cell-free expression system will contribute to the reduction in gene expression noise.

**Figure 4.**
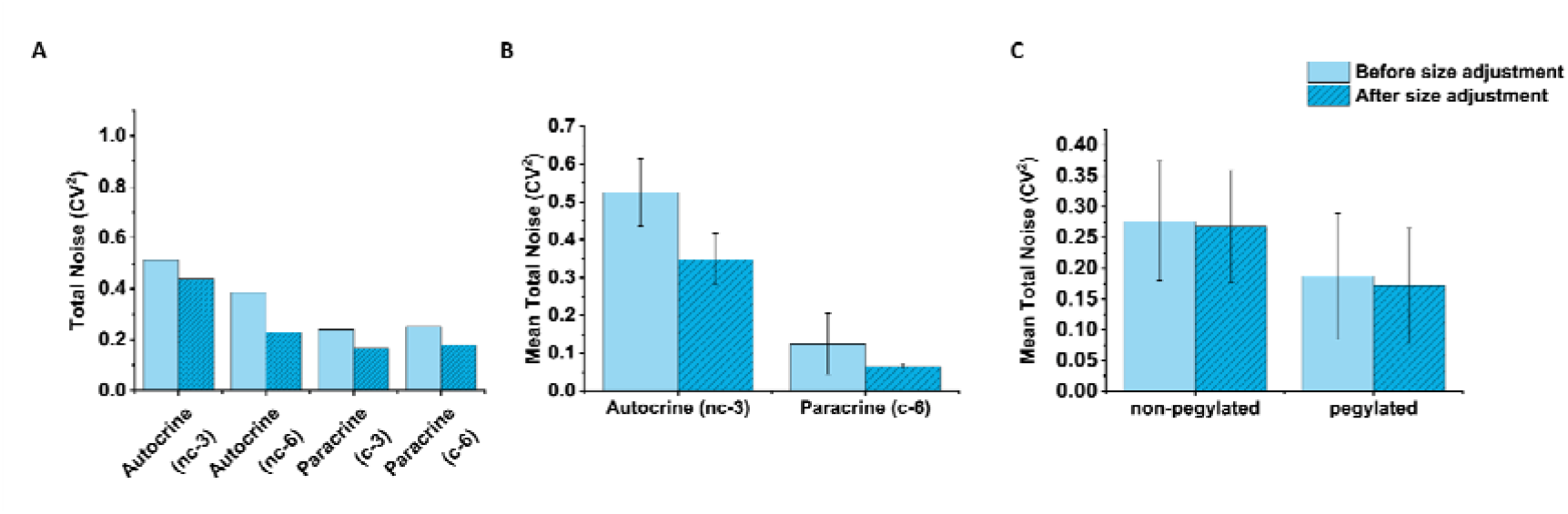
Size correction, communication architecture, and membrane pegylation reduce gene-expression variability in synthetic-cell populations. **(A)** Total expression noise (CV^2^) before and after vesicle-size adjustment across four circuit architectures: autocrine three-component, autocrine six-component, paracrine three-component, and paracrine six-component. Size adjustment reduced apparent variability in all conditions, indicating that vesicle diameter contributes to population-level fluorescence heterogeneity. **(B)** Mean total noise comparison between autocrine and paracrine architectures, showing lower variability in the communicating paracrine system compared with the non-communicating autocrine circuit. **(C)** Comparison of non-pegylated and pegylated vesicles before and after size adjustment. Pegylated vesicles showed reduced total expression noise, suggesting that membrane composition and vesicle–environment interactions contribute to reporter variability. Error bars indicate variation across replicate measurements, n = 4.

## Discussion

In this study we use cell-free gene circuits based on the Lux and Las systems from quorum sensing bacteria to define the parameters that can reduce population expression variability. The gene circuits were incorporated into lipids vesicles with cell-free expression systems using our on-chip inverted emulsion method. This method successfully allowed segregation of the circuits while allowing seamless diffusion of homoserine lactone signaling molecules^31^ between the vesicles using a chambered channel device. Analysis of gene expression output in individual vesicles across the population provided a route to compare the population variability (CV^2^) between autocrine and paracrine communication for three and six plasmid gene circuits. This provided a route to determine whether communication could reduce gene expression noise. Critically, our results show that contributing factors for the reduction in population variability are communication (Figure 1-2 and Supplementary figure 15), limitation of resources^32^ (Figure 3) and surface interactions^34^ (Figure 4). This could lead to approximately 60% reduction in noise. Identifying the precise mechanism or separating competing noise reduction contributions by which noise is reduced is still a challenge. However, the synthetic multi-cellular system with connected communication channels developed in this work is an ideal platform for design, build, test, and learn (DBTL) cycle framework. Following this, future work could investigate how genetic level changes, beyond plasmid concentration, like switching off of specific gene pathways or altering promoter strength can contribute to the noise reduction in synthetic multi-cellular systems. Overall, our work provides simple strategies to study and reduce gene expression variability to increase robustness in synthetic multi-cellular systems. This is important for using cell-free gene circuits for applications where robustness will be an important factor, for example for the design of collective systems for sensing, response and adapt to chemical cues and perform cell-free biological applications.

## Methods (See also Supplementary methods)

### Preparation of lipid vesicles using the inverted emulsion method

400 µM concentration of any given lipid composition (POPC 99.5% with 0.5% DID and when pegylated, 98.5%POPC, 0.5%DID and 1% DSPE-PEG 2000), dissolved in chloroform, was dried under a nitrogen stream and then under low pressure in a vacuum desiccator (45 min) to completely remove the solvent. The dried lipids were resuspended in 2 mL of mineral oil and sonicated in a water bath for 15 minutes to create a lipid dispersion (400 µM). The dispersion was sealed and stored at 4 °C until use. Next, chambered slides (Ibidi GmbH) were coated with a 2 mg/mL solution of β-casein to prevent non-specific binding between the glass slides and lipid vesicles. To do this, 100 µL of solution was loaded into the chamber and then centrifuged using Eppendorf 5430 R with swinging buckets at 900 ×g for 3 minutes to ensure uniform coverage and removal of any bubbles in the solution. The slides were then incubated for at least 2 hours in a desiccator until the β-casein coating is dry. The casein-coated slides are rinsed with 100 µL of an outer solution (Table S2b). An additional 100 µL of the outer solution is then added, followed by the addition of 25 µL of 400 µM lipid-oil dispersion to one chamber. In the case of uni-directional experiments, 25 µL of outer solution was added to the second chamber. For bi-directional experiments, 25 µL of 400 µM lipid-oil dispersion was added to the second chamber. The slide was incubated, at room temperature (23 °C) for 30 minutes to facilitate lipid monolayer formation between the oil and outer solution. During the wait time, a water-in-oil (W/O) emulsion was prepared by adding 5 µL of inner solution (Supplementary Table S2a) (Cell-free transcription– translation extract, energy mix, and the Lux/Las gene circuit) (see below table for specific plasmid types per condition) to 200 µL of the lipid-oil dispersion in a 1.5 mL Eppendorf tube. Droplets are typically formed by ratcheting the tube against Eppendorf tube rack/stand. 25 µL of the water-oil emulsion droplets were introduced to the chamber slide by pipetting into the chamber containing the lipid monolayer. One inlet in the case of one directional experiment and two in the case of the bidirectional experiments, each containing water-oil emulsions which contain different plasmids (see Supplementary Figure 2). Centrifugation was then performed at 3000 × g for 3 min which drives the droplets to the interface and facilitates the formation of lipid vesicles containing cell free expression system and plasmids in one or both the wells.

### Microscopy

Confocal microscopy of the vesicles was performed within the chambered slide using a system equipped with an HC PL APO 63x/1.30 GLYC CORR CS2 objective (Leica SP8) or S Plan Fluor 40x/0.60 objective (Nikon Eclipse Ti2). For imaging fluorescent labels such as eGFP, a 488 nm excitation wavelength was used, and emitted light was collected with filters centered at 520 ± 20 nm. In the case of mScarlet-I, 561 nm excitation was used, and emitted light was collected with filters centered at 580 ± 20 nm. For imaging membranes labeled with the either DiD dye (Ex, a 637 nm, Em collected at excitation wavelength was used, with corresponding filters at 650 ± 20 nm) or DHPE-Texas Red (Ex 565 nm, Em collected at 610 ± 20 nm). Time-lapse data was acquired at a resolution of 512 × 512 pixels with an interval of 5 minutes between frames at 30 °C (Temperature controller, H201-T-UNIT-BL, Okolab).

### Image analysis

Fluorescence microscopy images and time-lapse videos were acquired using the microscope software and subsequently analyzed in ImageJ/Fiji. Vesicle diameters and fluorescence intensities were measured manually from the acquired images. For each vesicle, the diameter was determined from the membrane channel, and reporter fluorescence intensity was measured from the corresponding eGFP and/or mScarlet-I channel. Vesicles that were out of focus, ruptured, overlapping, partially outside the field of view, or not clearly identifiable were excluded from the analysis. Time-lapse videos were used to follow changes in fluorescence intensity over time, and endpoint measurements were taken from the plateau phase of the fluorescence trajectory. No automated segmentation, machine-learning-based detection, or additional image-processing pipeline was used.

### Data analysis and fitting

All data analysis, statistical calculations, plotting, and curve fitting were performed using OriginPro software (OriginLab). Diameter and fluorescence intensity values obtained from ImageJ/Fiji measurements were exported and analyzed in OriginPro. For each condition, single-vesicle fluorescence intensities were used to calculate population statistics, including mean fluorescence intensity, standard deviation, coefficient of variation, and total noise. Total noise was calculated as the squared coefficient of variation,

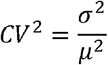

where µ is the mean single-vesicle fluorescence intensity and δ^2^ is the variance of the fluorescence intensity distribution. For time-lapse experiments, fluorescence intensities were averaged across vesicles at each time point to generate population-mean expression trajectories. Where indicated, fluorescence values were normalized to the maximum or plateau intensity of the corresponding trajectory to enable comparison between conditions. Kinetic traces were plotted and fitted in OriginPro using a four-parameter Boltzmann function,

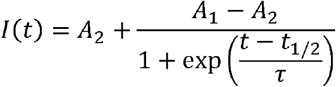

where A_1_ and A_2_ are the lower and upper fluorescence limits, t_1/2_ is the half-activation time, and τ is the characteristic time constant describing the steepness of the fluorescence increase. Endpoint fluorescence values were taken from the plateau phase of each trajectory. When different circuit architectures reached plateau at different times, the endpoint was selected from the time point at which the fluorescence intensity no longer changed appreciably with time. For selected comparisons, noise values were also calculated from three or more sequential subsets of the data, and the mean CV^2^ and standard error of the mean were reported.

### Size dependency estimation

To account for vesicle-size-dependent expression, fluorescence intensity was corrected for its dependence on vesicle diameter. For each condition, endpoint fluorescence intensity was plotted as a function of vesicle diameter, and the size dependence was fitted by linear regression. *I*_*i*_ = *β*_0_ + *β*_1_ *D*_*i*_ + *ε*_*i*_ where I_i_ is the fluorescence intensity of vesicle (i), D_i_ is its diameter, β_0_ and β_1_ are the fitted regression parameters, and ε_i_ is the residual fluorescence not explained by vesicle size. The expected fluorescence intensity based on vesicle diameter was calculated as: *I*_*fit,i*_ = *β*_0_ + *β*_1_ *D*_*i*_ and the residual was calculated as: *ε*_*i*_ = *I*_*i*_ − *I*_*fit,i*_. Size-adjusted fluorescence values were then obtained by adding the residuals back to the population mean fluorescence: *I*_*adjusted,i*_ = *µ* + *ε*_*i*_ where µ is the mean fluorescence intensity of the unadjusted population. Size-adjusted total noise was calculated from these corrected fluorescence values (*I*_*adjusted,i*_):

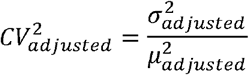

This correction retained all vesicles and removed the systematic fluorescence variation associated with vesicle diameter, rather than selecting vesicles within a restricted size range. For the case of Autocrine (nc-6), a single-exponential decay model, I=y0 +Ae−D/t is used.

### Gillespie stochastic simulation algorithm (SSA) based Plasmid Copy Number simulations

Stochastic gene-expression dynamics were simulated for four circuit architectures: non-communicating synthetic cells (SCs), uni-directional SC communication, all-in-one systems, and bidirectional communication, each derived from our LUX/LAS quorum-sensing designs based on [ref]. Simulations were performed using a custom Gillespie stochastic simulation algorithm (SSA) implementation (Python 3.10 with NumPy, pandas, and Matplotlib). All kinetic parameters were fixed across sweeps:

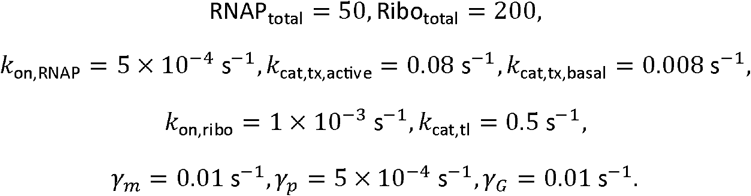

This parameter set defines a baseline operating regime following the resource-competition modeling strategy of Goetz *et al*. (2025). The values of RNAP_total_ = 50, Ribo_total_ = 200, and the associated binding, transcription, translation, maturation, and degradation rates represent an effective coarse-grained parameterization of the system rather than direct biochemical measurements. Each plasmid copy-number condition (5, 10, 20, 40) was simulated using *N* = 500SSA trajectories up to *T*_end_ = 2000 s. Trajectories were sampled at integer-second resolution, and the steady-state window was defined as *t* = 1500−2000 s. For communicating architectures, producer and responder compartments exchanged AHL (and Las signal for bidirectional communication) via first-order inter-compartment transport with mean travel time of 5 secs corresponding to a 1 cm separation.

## Data Availability

Source data for the Figures is available here - https://doi.org/10.6084/m9.figshare.33973018

## Code availability

The Python code used to perform the Gillespie stochastic simulation algorithm (SSA) plasmid copy-number sweeps and RNAP/ribosome resource sweeps is available on GitHub at: https://github.com/nyandrapalli/Plasmid-and-Resource-sweeps. The repository contains separate folders for the plasmid sweep and resource sweep simulations, including model scripts, and sweep runners, used to generate the simulation results reported in this study.

## Acknowledgements

Funding was provided by the European Union (European Research Council [ERC], MinSynCell, 101088834., and ITN, DARCHEMDN grant number 101119956, awarded to T.-Y.D.T.). The views and opinions expressed are, however, those of the authors only and do not necessarily reflect those of the European Union or the ERC; neither the European Union nor the granting authority can be held responsible for them. We acknowledge financial support from the University of Saarland and the Max Planck Society (T.-Y.D.T.). NY. and S.S. were funded through ERC, MinSynCell, 101088834. We thank the PharmaScienceHub, Saarland University. for financial support (T.-Y.D.T.). We thank the Light Microscopy Facility (LMF), Advanced Imaging Facility (AIF), of the Max Planck Institute of Molecular Cell Biology and Genetics (MPI-CBG) for their technical support and useful discussions. We thank the Advanced Light Microscopy Facility at the Saarland University. We thank Christoph Zechner for fruitful discussion regarding this work. We acknowledge contributions from Surased Suraritdechachai and Anjali Anilkumar for support in preparation of materials.

## Author Contributions

T-Y DT and DTG conceived the study, TY-DT, NY and DG designed the experiments; NY undertook the experiments. TYDT, NY, DG wrote the manuscript. All authors contributed approved the text.

