## Supplementary information for "Gene expression noise is reduced in communicating synthetic cell populations"

Materials and methods:

| REAGENT or RESOURCE | SOURCE | IDENTIFIER |
| --- | --- | --- |
| *Chemicals* |  |  |
| L-Alanine | Sigma, USA | A7627 |
| L-Arginine | Sigma, USA | A5006 |
| L-Asparagine | Sigma, USA | A0884 |
| L-Aspartic acid | Sigma, USA | A9256 |
| L-Cysteine | Sigma, USA | W326305 |
| L-Glutamic acid | Sigma, USA | G1251 |
| L-Glutamine | Sigma, USA | G3126 |
| Glycine | Sigma, USA | G7126 |
| L-Histidine | Sigma, USA | H8000 |
| L-Isoleucine | Sigma, USA | I2752 |
| L-Leucine | Sigma, USA | L8000 |
| L-Lysine | Sigma, USA | L5501 |
| L-Methionine | Sigma, USA | M9625 |
| L-Phenylalanine | Sigma, USA | P2126 |
| L-Proline | Sigma, USA | P0380 |
| L-Serine | Sigma, USA | S4500 |
| L-Threonine | Sigma, USA | T8625 |
| L-Tryptophan | Sigma, USA | T0254 |
| L-Tyrosine | Sigma, USA | T3754 |
| L-Valine | Sigma, USA | V0500 |
| Sucrose | Sigma, USA | 1076539053 |
| Glucose | Sigma, USA | G8270 |
| NTPs |  |  |
| Adenosine 5′-triphosphate disodium salt hydrate | Sigma, USA | A26209 |
| Cytidine 5′-triphosphate disodium salt hydrate | Sigma, USA | 30320 |
| Guanosine 5′-triphosphate sodium salt hydrate | Roche, CH | 10106399001 |
| Uridine 5′-triphosphate trisodium salt dihydrate | Sigma, USA | 94370 |
| *Cofactors* |  |  |
| β- Nicotinamide adenine dinucleotide (NAD) | Sigma, USA | N1511 |
| Coenzyme A (CoA) | Sigma, USA | C4284 |
| Folinic acid calcium salt hydrate | Sigma, USA | F7878 |
| Oxalic acid | Roth, DE | 8879.1 |
| Phosphoenolpyruvate (PEP) | Roche, CH | 10108294001 |
| Putrescine | Sigma, USA | 51799 |
| Spermidine | Sigma, USA | Sigma, USA |
| tRNA from E. coli MRE600 | Roche, CH | 10109541001 |
| *Materials for Buffers* |  |  |
| Acetic acid (HOAc) | Merck, USA | K48001663 632 |
| Dithiothreitol (DTT) | Thermo, USA | R0862 |
| HEPES (N-2-Hydroxyethylpiperazine-N'-2- ethane sulphonic acid) | Carl Roth, DE | 9105 |
| L-Glutamic acid hemimagnesium salt tetrahydrate | Sigma, USA | 49605 |
| L-Glutamic acid potassium salt monohydrate | Sigma, USA | G1149 |
| Potassium hydroxide | Sigma, USA | 221473 |
| Potassium phosphate dibasic solution | Sigma, USA | P8584 |
| Potassium phosphate monobasic solution | Sigma, USA | P8709 |
| Trizma base | Sigma, USA | T1503 |
| *Materials for Oil phase* |  |  |
| Mineral Oil | Sigma, USA | M5904 |
| *Materials for Lipid phase* |  |  |
| 1,2-dioleoyl-sn-glycero-3-phosphocholine (DOPC) | Avanti, USA | 850356C |
| Biotin PE | Avanti, USA | 860562C |
| 1-palmitoyl-2-oleoyl-sn-glycero-3-phosphocholine | Avanti, USA | A80557 |
| 1,2-distearoyl-sn-glycero-3-phosphoethanolamine-N-[amino(polyethylene glycol)-2000] (ammonium salt) | Avanti, USA | A88128 |
| 1,1'-dioctadecyl-3,3,3',3'-tetramethylindodicarbocyanine, 4-chlorobenzenesulfonate salt (DiD) | ThermoFisher, USA | D7757 |
| 1,2-Dihexadecanoyl-sn-Glycerin-3-Phosphoethanolamin, Triethylammoniumsalz (DHPE Teax Red) | ThermoFisher, USA | T1395MP |
| Chambered slides (µ-Slide VI 0.5 Glass Bottom | Ibidi | 80607 |

Bacterial strain list:

| Strain name | Description | Source |
| --- | --- | --- |
| E. coli DH5α | High efficiency competent cells for transformation. | NEB (C2987) |
| E. coli BL21 (DE3) | Competent cells for T7 RNAP-mediated expression. | NEB (C2527) |

Kits list:

| Name | Supplier | Catalog NO. |
| --- | --- | --- |
| NEBuilder HiFi DNA Assembly Kit | NEB, USA | E2621 |
| QIAprep Spin Miniprep Kit | QIAGEN, DE | 27104 |
| QIAGEN Plasmid Maxi Kit | QIAGEN, DE | 12162 |
| QIAquick PCR Purification Kit | QIAGEN, DE | 28104 |
| myTXTL Sigma 70 Master Mix Kit | Daicel Arbor Biosciences, USA | 507024 |

Equipment list:

| Name | Supplier | Catalog No. |
| --- | --- | --- |
| Spectra/Por 2 Dialysis Membrane (12-14 kD) | Repligen, USA | 132680T |
| Avanti Centrifuge J26-XP with JLA-8.1000 rotor | Beckman Coulter, USA | 363688 |
| NanoDrop 2000 | Thermo, USA | ND-2000 |
| Sonorex sonication bath | Bandelin, DE | - |
| HC PL APO **63x**/1,30 GLYC CORR CS2 | Leica microsystems, DE | 11506353 |
| S Plan Fluor 40x/0.60 objective | Nikon Eclipse Ti2 | 88-381 |
| Osmomat 3000 | Gonotec | - |
| Centrifuge 5430 R with swinging titre plate rotor | Eppendorf | EP5428000010 |

Plasmids: All plasmids used this study have been previously published.

NT/pT7 LasI (Addgene #193631), pEXP5-NT/pLaseGFP (Addgene #193630), pEXP5-NT/pT7 LasR (Addgene #193629), pEXP5-NT/pLux LasI (Addgene #193628), pEXP5-NT/pLuxLuxI (Addgene #193627), pEXP5NT/pT7 LuxI (Addgene #193626), pEXP5-NT/pT7 LuxR (Addgene #193625), pEXP5-NT/pLuxeGFP (Addgene #193624)  pEXP5-NT/eGFP plasmid (<https://doi.org/10.1039/c1sc00183c>).

The plasmid assembly of pEXP5-NT/pLas mScarletI was performed by using NEBuilder HiFi DNA assembly mastermix (NEB, USA). The mScarlet gene was amplified using the following primers:GCAGCGGCGAAAACCTGTATTTTCAGTCCATGGTTTCAAAAGGAGAAGC and GGCTTTGTTAGCAGCCGGATCACCCTTTACTTGTACAGTTCATCCATACC. The accepting vector fragments were amplified from pEXP5-NT/pLas eGFP with two primer pairs: TAAAGGGTGATCCGGCTGC and TAGATAACTACGATACGGGAGG, TATCATTGCAGCACTGGG and GGACTGAAAATACAGGTTTTCG. The plasmid was transformed into *E. coli* DH5α. All plasmids were prepared from *E. coli* DH5α using QIAGEN Plasmid Maxi Kit (QIAGEN, Germany) as described by the manufacturer.

Preparation of cell free extract

The process for preparing E. coli cell-free extract was prepared as previously described [ref]. In brief, the process starts with an overnight starter culture of E. coli BL21 (DE3) cells grown in LB media. This is then used to inoculate two 500 mL pre-warmed cultures of 2xYTP media (1 L of a solution containing 5 g NaCl, 10 g yeast extract, 16 g tryptone, 40 mL of 1 M potassium phosphate dibasic solution, and 22 mL of 1 M potassium phosphate monobasic solution. The final volume was adjusted to 1 L with water), which are incubated until an optical density of 1.6 is reached. The cells are then harvested via centrifugation, and the resulting pellet is washed three times with S30A buffer (500 mL of a buffer with 14 mM MgGlu, 60 mM KGlu, 50 mM Tris, and 2 mM DTT, adjusted to pH 7.7 using approximately 1 mL of glacial acetic acid). The wet cell pellet is weighed, flash-frozen in liquid nitrogen, and stored overnight at -80 °C. The following day, the frozen pellet is thawed and resuspended in S30A buffer at a ratio of 1 mL of buffer per 1 g of wet pellet. The resuspended cell solution is aliquoted, and the cells are lysed through sonication in an ice-water bath using a pulsatile method to prevent overheating. Immediately after sonication, DTT is added to each aliquot. The lysate undergoes two clarification steps via centrifugation to remove cellular debris. The first clarification is followed by a "run-off reaction" where the supernatant is incubated at 37 °C for one hour to degrade unwanted nucleic acids and proteins. After the second clarification, the lysate is dialyzed against S30B buffer (1 L of a buffer containing 14 mM MgGlu, 60 mM KGlu, and 2 mM DTT, adjusted to pH 8.2 with about 2 mL of 2 M Tris.) for three hours. The final dialyzed lysate is subjected to one last clarification step (the dialyzed lysate was centrifuged at 10,000 xg for 10 min), aliquoted into 50 μL volumes, flash-frozen in liquid nitrogen, and stored at -80 °C for subsequent use.

Preparation of energy and nutrient mixture for gene expression:

**Solution A**

The stock and final concentrations of the components used for Solution A are listed in Table S7. All stock solutions were prepared by dissolving in water and the HEPES buffer was adjusted to pH 7 using KOH. 50 μL aliquots of Solution A were flash frozen with liquid nitrogen and stored at -80°C until use.

**Solution B**

The stock and final concentrations of the components of Solution B are listed in Table S8. The amino acid mix contains 50 mM of each amino acid and was incubated at 37°C with shaking to help dissolve the powder. Note that the amino acids may not dissolve completely in solution. The amino acid mix is prepared in a large volume of 10 mL to ensure that sufficient material could be weighed out. The amino acid solution should be fully mixed before adding into the final Solution B mixture. The stock of magnesium glutamate and potassium glutamate were prepared as one solution. The PEP stock solution was adjusted to pH 7 using 10 M KOH solution. The amounts of the final Solution B mix are given to make 2.1 mL of Solution B or approximately 1000 reactions. 50 μL aliquots of Solution B were flash frozen with liquid nitrogen and stored at -80°C until use.

**Table S1.** List of reagents used in Solution A and Solution B

| **Solution A** | | **Solution B** | |
| --- | --- | --- | --- |
| Component | Final (mM) | Component | Final (mM) |
| ATP | 12.4 | Amino acid mix | 14.3 |
| GTP | 8.7 | Magnesium glutamate | 71 |
| CTP | 8.7 | Potassium glutamate | 930 |
| UTP | 8.7 | PEP (pH 7) | 238 |
| Folinic acid | 0.68 |  |  |
| tRNA | 0.176 mg/mL |  |  |
| NAD | 2.7 |  |  |
| CoA | 1.8 |  |  |
| Oxalic acid | 27.2 |  |  |
| Putrescine | 6.8 |  |  |
| Spermidine | 10.1 |  |  |
| HEPES (pH 7) | 774.5 |  |  |

**The home-made CFES reaction mixture:**

The final E. coli extract-based CFES master mix is prepared according to Table S2 and incubated at 30°C for 5-10 hours. The amount of magnesium glutamate supplemented to the final CFES optimized to be 2mM. To prepare the CFES mix, the DNA + water components of each sample were first pre-mixed, the remainder of the master mix was then added to avoid long waiting times between starting the CFE reactions. All encapsulated experiments in this study follow the CFES mix.

**Table S2a.** The home-made CFES mixture (inner solution) for synthetic cell preparation

| **Component** | **µL** |
| --- | --- |
| Solution A | 1.8 |
| Solution B | 1.75 |
| bCFES | 5 |
| MgGlu (200 mM) | 0.156 |
| Plasmids | 0.625 × (number of plasmid types) |
| Ribose (30 mM) | 0.625 |
| Maltodextrin (60 mM) | 0.625 |
| Nuclease-free Water | 4 – (0.625 × number of plasmid types) |

**Table S2b.** The outer solution for synthetic cell preparation (multiplied to desired volume)

| **Component** | **µL** |
| --- | --- |
| Solution A | 1.8 |
| Solution B | 1.75 |
| Nuclease-free Water | 6.25 |
| MgGlu (200 mM) | 0.156 |
| Ribose (30 mM) | 0.625 |
| Maltodextrin (60 mM) | 0.625 |

**Table S3**: List of various combinations of plasmids used within lipid vesicles:

|  | Conditions | Encapsulated Plasmid types | |
| --- | --- | --- | --- |
|  |  | SCs Population 1 | SCs Population 2 |
| (i) | Non-communicating SCs / Autocrine (nc-3) | pT7-LUX I + pT7 LUX R + pLUX GFP | NA |
| (ii) | Unidirectional SCs / Paracrine (c-3) | pT7-LUX I | pT7-LUX R  +  pLUX GFP |
| (iii) | Bidirectional SCs / Paracrine (c-6) | pT7-LUX I + pT7-LAS R + pLAS GFP | pLUX LAS I +  LUX R  + pLUX GFP |
| (iv) | All-in-one / Autocrine (c-6) | pT7 LUX I + pT7LAS R + pLAS mScarlet +  pLUX LAS I +  pT7 LUX R  + pLUX GFP | NA |
| (v) | Control | pT7 eGFP | NA |

**Supplementary Methods – Plasmid Copy Number Sweeps**

**Model Overview**

Each condition is simulated with an exact Gillespie SSA implementation over integer time steps.

**Stochastic formulation and finite-resource coupling:**

The Gillespie stochastic simulation algorithm (SSA) determines both the waiting time to the next reaction and the identity of that reaction. For a system in state $x$, each reaction channel $j$has a propensity $a_{j}(x)$, and the total propensity is:

$$a_{0}(x)=\sum_{j} a_{j}(x)$$

The waiting time $\tau$until the next reaction is drawn from an exponential distribution with rate $a_{0}(x)$:

$$\tau=-\frac{\ln(r_{1})}{a_{0}(x)}$$

The probability that reaction $j$fires next is:

$$P(\text{next reaction}=j\mid x)=\frac{a_{j}(x)}{a_{0}(x)}$$

Mass‑action propensities:

$$a_{\text{bind,RNAP}}=k_{\text{on,RNAP}}\text{ }\text{RNAP}_{\text{free}}\text{ }P_{\text{free}}$$

$$a_{\text{tx,active}}=k_{\text{cat,tx,active}}\text{ }P_{\text{RNAP}}$$

$$a_{\text{tx,basal}}=k_{\text{cat,tx,basal}}\text{ }P_{\text{RNAP}}$$

$$a_{\text{bind,ribo}}=k_{\text{on,ribo}}\text{ }\text{Ribo}_{\text{free}}\text{ }m$$

$$a_{\text{tl}}=k_{\text{cat,tl}}\text{ }R_{\text{bound}}$$

$$a_{\text{deg,m}}=\gamma_{m}\text{ }m$$

$$a_{\text{deg,p}}=\gamma_{p}\text{ }p$$

$$a_{\text{mat}}=k_{\text{mat}}\text{ }G_{i}$$

$$a_{\text{syn,AHL}}=k_{I}\text{ LuxI}$$

$$a_{\text{syn,S}}=k_{S}\text{ LasI}$$

Finite‑resource constraints:

$$\text{RNAP}_{\text{free}}+\sum(\text{RNAP–promoter complexes})=\text{RNAP}_{\text{total}}$$

$$\text{Ribo}_{\text{free}}+\sum(\text{ribosome–transcript complexes})=\text{Ribo}_{\text{total}}$$

Inter‑compartment diffusion:

$$a_{A\to B}=k_{\text{diff}}\text{ }X_{A}$$

$$a_{B\to A}=k_{\text{diff}}\text{ }X_{B}$$

For each trajectory set we computed GFP means, standard deviations, and coefficient of variation squared (CV² = (SD/mean)²) over the steady-state window. Three independent sweeps (distinct random seeds) were generated per condition and copy number; the manuscript reports the mean and standard deviation of CV² across these repeats. All simulation scripts are provided in the accompanying GitHub repository (<https://github.com/nyandrapalli/plasmid-copy-sweeps/tree/master>).

**Table S4.** Circuit configurations used in the plasmid copy-number sweeps, summarizing promoter/species composition and signaling assumptions for each model.

| **Condition** | **Circuit Description** |
| --- | --- |
| Cond1 | Single “producer + reporter” cell carrying pT7–luxI and pLux–GFP (LuxR constitutive). |
| Cond4 | Single cell containing the full Cond3 plasmid set (pT7–luxI/lasR/luxR, pLas–GFP/mScarlet, pLux–GFP, pLux–lasI). |
| Cond5 | Two-cell circuit with finite-distance LuxI producer and LuxR/pLux–GFP responder that share AHL via first-order exchange (distance = 1 cm, mean travel time = 5 s). |
| Cond6 | Two-cell Lux/Las cross-talk; both AHL and Las signals diffuse between the compartments with the same 5 s travel time at 1 cm separation. |

Shared kinetic parameters: RNAP_total = 50, Ribo_total = 200, kon_act = 1×10⁻³ s⁻¹, koff_act = 1×10⁻³ s⁻¹, kon_RNAP = 5×10⁻⁴ s⁻¹, kcat_tx_active = 0.08 s⁻¹, kcat_tx_basal = 0.008 s⁻¹, kon_ribo = 1×10⁻³ s⁻¹, kcat_tl = 0.5 s⁻¹, γ_m = 0.01 s⁻¹, γ_p = 5×10⁻⁴ s⁻¹, γ_G = 0.01 s⁻¹.

Shared kinetic parameters: $\text{RNAP}_{\text{total}}=50,\text{ }\text{Ribo}_{\text{total}}=200,\text{ }k_{\text{on,act}}=1\times{10}^{-3}\text{ }\text{s}^{-1},\text{ }k_{\text{off,act}}=1\times{10}^{-3}\text{ }\text{s}^{-1},\text{ }k_{\text{on,RNAP}}=5\times{10}^{-4}\text{ }\text{s}^{-1},\text{ }k_{\text{cat,tx,active}}=0.08\text{ }\text{s}^{-1},\text{ }k_{\text{cat,tx,basal}}=0.008\text{ }\text{s}^{-1},\text{ }k_{\text{on,ribo}}=1\times{10}^{-3}\text{ }\text{s}^{-1},\text{ }k_{\text{cat,tl}}=0.5\text{ }\text{s}^{-1},\text{ }\gamma_{m}=0.01\text{ }\text{s}^{-1},\text{ }\gamma_{p}=5\times{10}^{-4}\text{ }\text{s}^{-1},\text{ }\gamma_{G}=0.01\text{ }\text{s}^{-1}.$

Promoter copy numbers were varied as described below.

**Simulation Protocol**

- **Plasmid copies:** {5, 10, 20, 40} plasmids per plasmid type; in Cond5/Cond6 both cells use the same copy number for all plasmids.
- **Replicates:** N = 500 trajectories per condition × copy combination (five batches of 100 trajectories to avoid runtime limits).
- **Temporal settings:** T_end = 2000 s; trajectories sampled at integer seconds. Steady-state window defined as 1500–2000 s.
- **Outputs per run:**
  - Raw GFP traces (CSV) with one column per time point (GLas_t0, GLux_t0, etc.).
  - Steady-window statistics (mean, SD, CV, CV², N) saved to condX_summary.csv.
  - Mean and percentile plots (condX_timelapse_plot.png) showing mean ± 10–90% envelopes.
- **Repeat strategy:** Three independent sweeps per condition and copy number (original run plus two repeats) were generated with different random seeds. CV² statistics reported in the manuscript are the mean ± SD across these three sweeps.

**Implementation**

**Table S5.** Scripts reside in scripts for plasmid copy number sweeps:

| **Script** | **Description** |
| --- | --- |
| cond1_plasmid_copy_sweep.py | Calls cond1_timelapse.run() for each copy. |
| cond4_plasmid_copy_sweep.py | Loads Trail sweep 1/repo/scripts/cond4_timelapse.py via importlib and sweeps copies. |
| cond5_plasmid_copy_sweep.py | Runs copy sweeps with distance = 1 cm, τ = 5 s diffusion. |
| cond6_plasmid_copy_sweep.py | Equivalent sweep for the cross-talk system (both AHL and Las diffusion). |

Each driver sets N=500, T_end=2000, window=(1500,2000), and writes results into Trails_plasmid/condX_plasmid_sweeps/copies_<copy>/.

**Reproducibility**

- **Environment:** Python 3.10 micromamba environment with NumPy, pandas, Matplotlib.
- **Command example:**
  micromamba run -p C:\Users\admin\micromamba_env python scriptsforplasmidcopynumbersweeps/cond6_plasmid_copy_sweep.py
- **Aggregation:** After each sweep, raw batches were merged (five × 100 trajectories) and steady-window statistics were recalculated to produce a single condX_summary.csv per copy. The reported CV² values are compiled in all_conditions_cv2_summary.csv.

Supplementary figures:

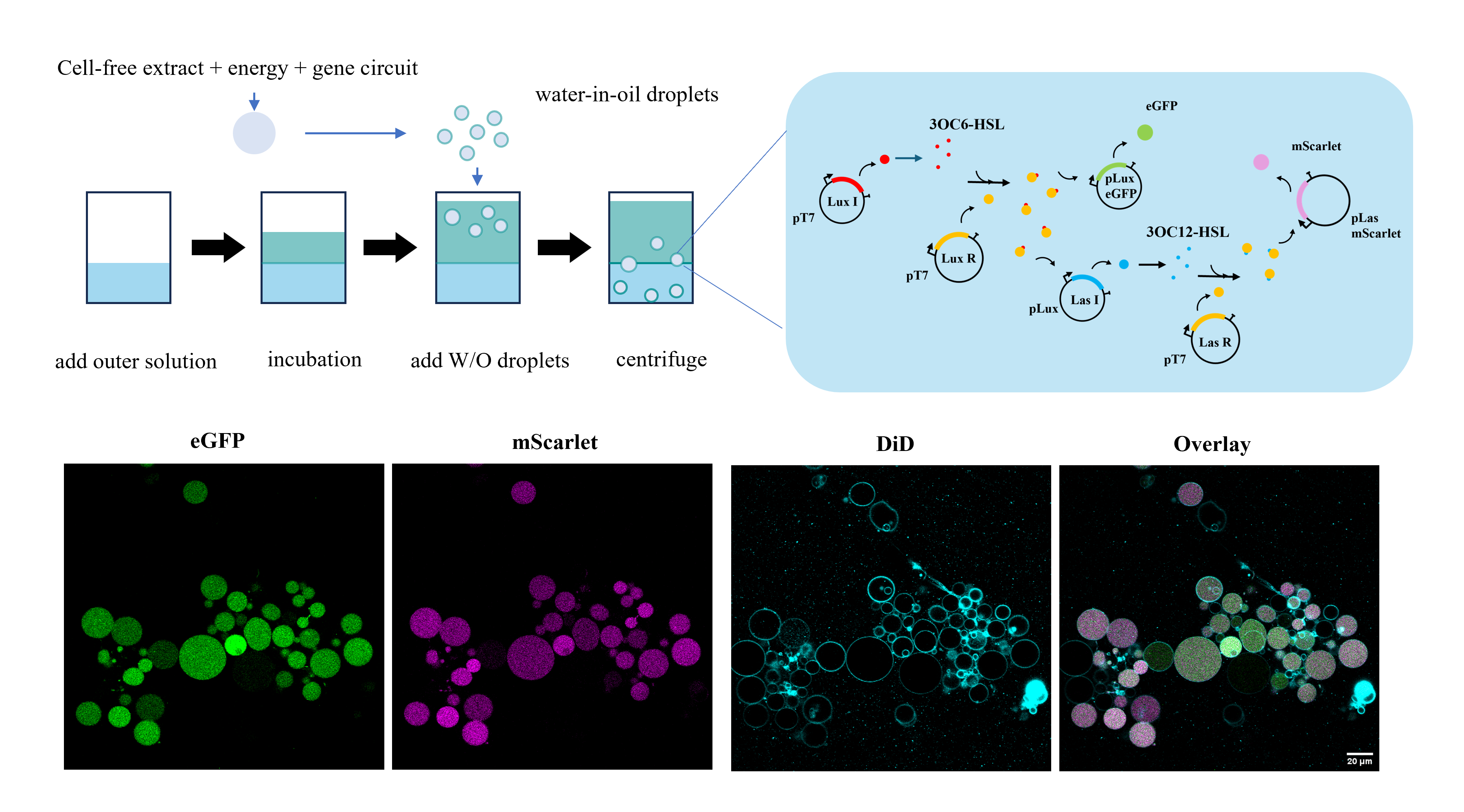

**Supplementary figure 1:** **Cell-free expression of quorum sensing gene circuit within SCs population**. Top: Schematic of the inverted emulsion protocol used to generate gene-expressing giant unilamellar vesicles (GUVs). Cell-free transcription–translation extract, energy mix, and the Lux/Las gene circuit are first encapsulated in water-in-oil droplets. These droplets are transferred across a lipid-stabilized interface by centrifugation to yield GUVs in an outer solution. Insert showing the Lux/Las gene circuitry: LuxI-driven AHL (N-Acyl Homoserine Lactone) production activates LuxR to induce LasI and the reporter eGFP, while LasI generates a second AHL signal that, together with LasR, activates mScarlet expression. Bottom: Confocal images of synthetic cells co-expressing eGFP and mScarlet. Bottom: eGFP (green, left) and mScarlet (magenta, right) fluorescence from the encapsulated gene circuit. Bottom left: DiD-labeled membranes outlining the vesicles. Bottom right: overlay of all channels, showing homogeneous co-localization of both reporters within membrane-bounded compartments. Scale bar, 20 µm.

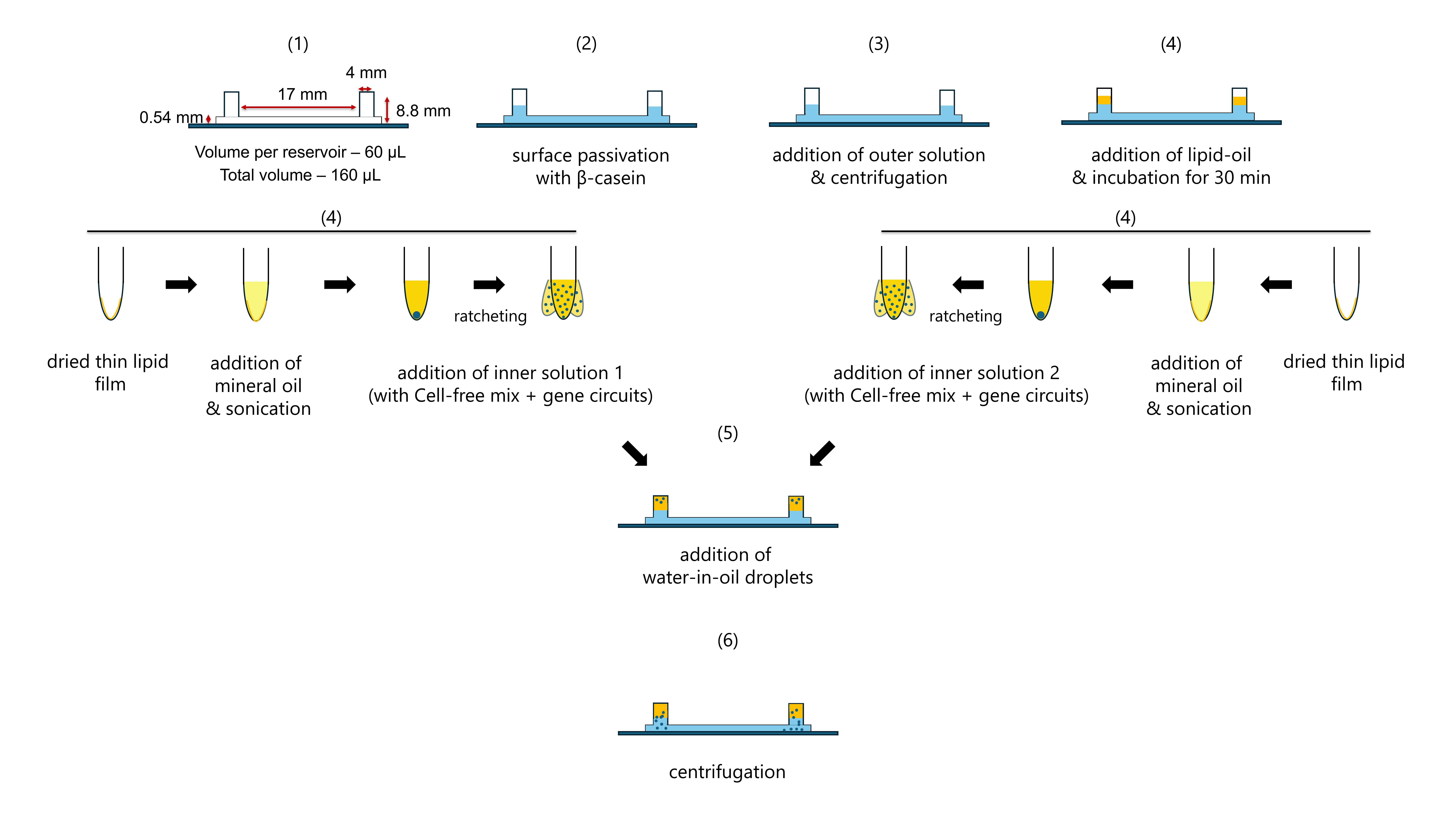

Supplementary figure 2. Preparation of phase-transfer chambers and water-in-oil droplets for synthetic cell assembly (1) Geometry of the two-reservoir observation chamber used for phase transfer (height 0.54 mm, reservoir length 17 mm, central lane width 4 mm; 60 µL per reservoir, total volume 160 µL). (2) The glass surface is passivated with β-casein. (3) Outer solution is added and equilibrated by centrifugation. (4) A lipid–oil mixture is layered on top and incubated for 30 min to form a stable lipid monolayer. In parallel, thin lipid films are rehydrated in mineral oil and sonicated, followed by addition of inner solution 1 or 2 containing cell-free extract and desired gene circuits; repeated pipetting (“ratcheting”) generates water-in-oil droplets. (5) The droplets are transferred into the prepared chamber with the monolayer. (6) Centrifugation drives the droplets across the lipid-stabilized interface, yielding two populations of giant lipid vesicles (synthetic cells) in the central observation lane.

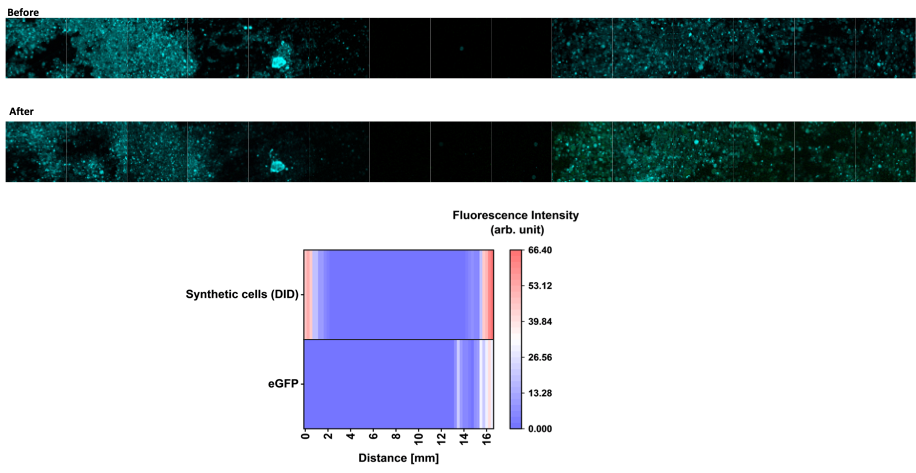

**Supplementary figure 3: Paracrine (c-3) cell populations visualization along the entire chip length.** Top: Confocal images of the Paracrine (c-3) communication lane before and after, showing the expression of eGFP only at the right side of the channel. Bottom: Heat map spatial profile of Synthetic cells and eGFP intensity along the communication lane at steady state, showing bright activation zone while the opposing SC population (left) is with no signal, similar to the central region.

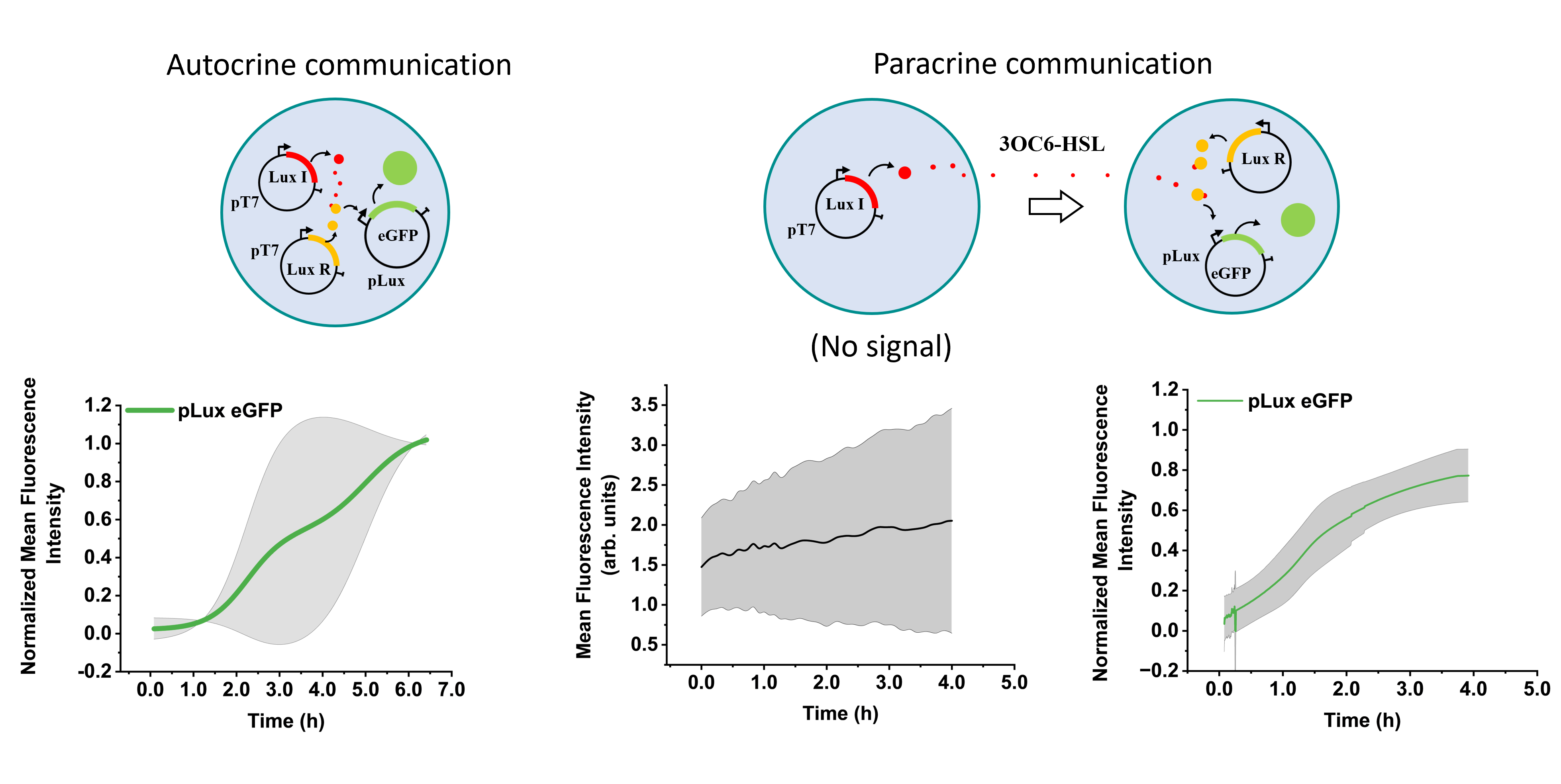

**Supplementary figure 4:** **Gene expression profile of Autocrine and Paracrine Lux only systems.** Left: Schematic showing the three plasmids, LuxI produces AHL that activates LuxR and drives eGFP expression. Population-averaged pLux–eGFP trajectory (green), showing a delayed but sigmoidal activation of the Autocrine (nc-3) population. Right: Schematic showing the three plasmids, separated across two synthetic cell populations, LuxI produces AHL diffuses out of the SCs and activates LuxR in the second population and drives eGFP expression. Population-averaged no expression flat line is observed for first population with only pT7 Lux-I, while the second population with pT7 Lux R and pLux–eGFP show increasing trajectory (green), showing sigmoidal activation of the Paracrine (c-3) communication.

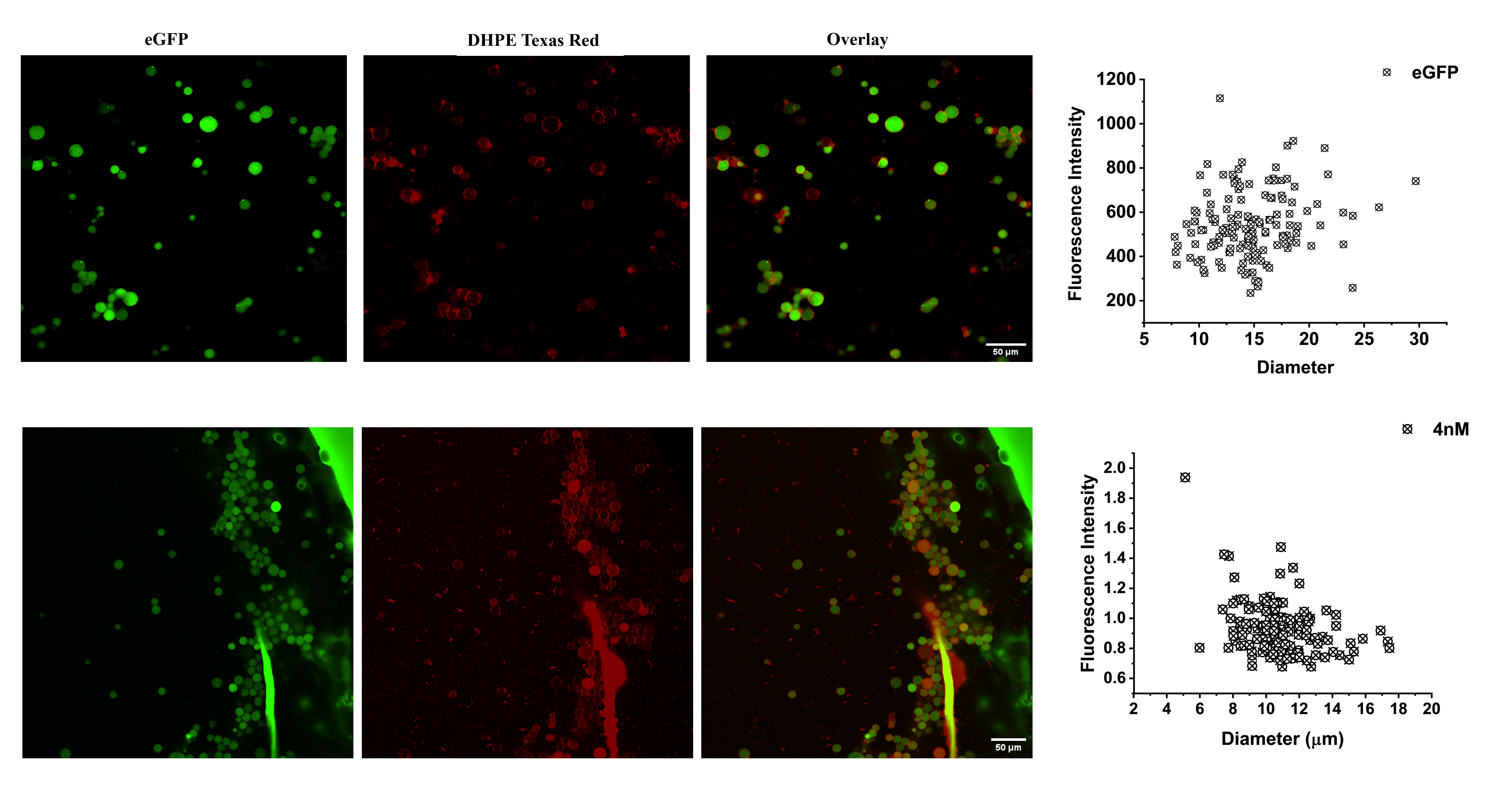

**Supplementary figure 5: eGFP reporter protein in SCs populations.**

Top (left to right): Confocal images showing encapsulated purified eGFP protein within SCs with eGFP in green, SCs labelled in red with DHPE-Texas Red. Overlay showing the localization of the protein within the SCs. Graph showing the fluorescence intensity of encapsulated eGFP protein within the SCs of varying diameter. Bottom (left to right): Confocal images showing gene expression of pT7 eGFP (4nM) within SCs with eGFP in green, SCs labelled in red with DHPE-Texas Red. Overlay showing the localization of the protein within SCs. Graph showing the fluorescence intensity of expressed eGFP protein within the SCs of varying diameter.

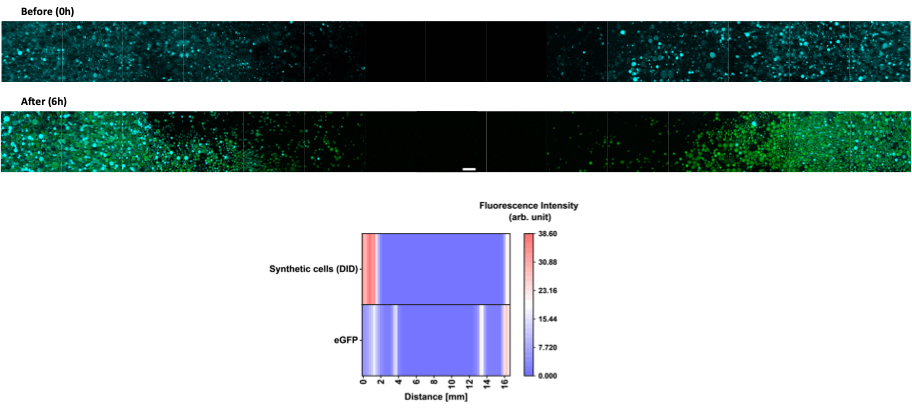

**Supplementary figure 6: Paracrine (c-6) cell populations visualization along the entire chip length.** Top: Confocal images of the Paracrine (c-6) communication lane before and after, showing the expression of eGFP on both sides of the channel. Bottom: Heat map spatial profile of Synthetic cells and eGFP intensity along the communication lane at steady state, showing bright activation zones for both the synthetic cell populations, with no signal at the central region where SCs are not present.

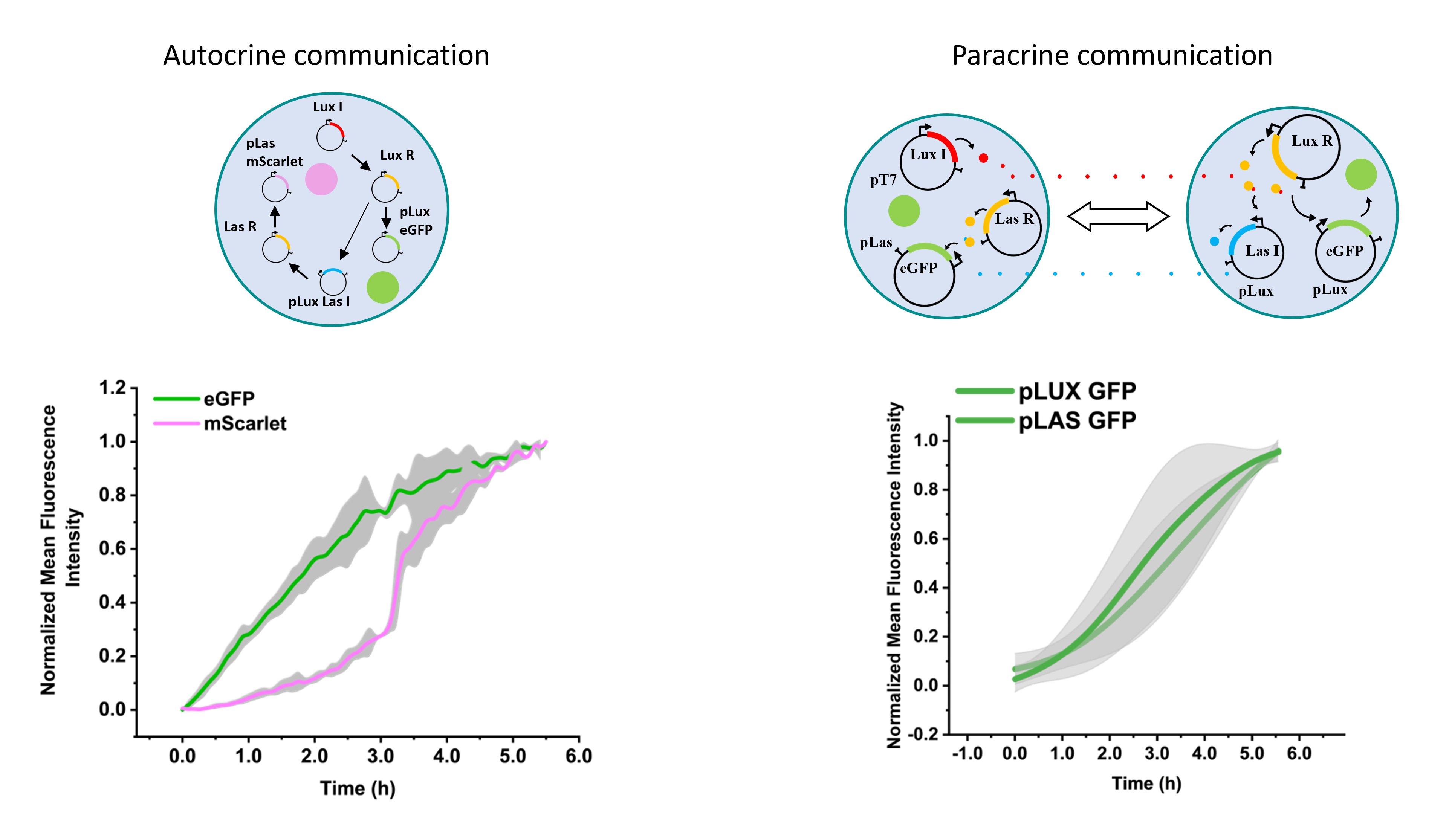

**Supplementary figure 7: Gene expression profile of Autocrine and Paracrine Lux/Las combined systems.** Left: Schematic showing the six plasmids, LuxI-driven AHL (N-Acyl Homoserine Lactone) production activates LuxR to induce LasI and the reporter eGFP, while LasI generates a second AHL signal that, together with LasR, activates mScarlet expression. Population-averaged pLux–eGFP expression trajectory (green), pLas–mScarlet expression trajectory showing sigmoidal activation of the Autocrine (nc-6) population. Right: Schematic showing the six plasmids, separated across two synthetic cell populations, LuxI produces AHL diffuses towards the second population of SCs, together with LuxR in the second population, driving eGFP expression. The AHL activates pLux LasI, inducing the production of AHL that diffuses out towards the LasR and pLux mScarlet containing SCs and inducing mScarlet production. Graph showing the increasing trajectories (green) from both the SC populations - showing sigmoidal activation of the Paracrine (c-6) communication.

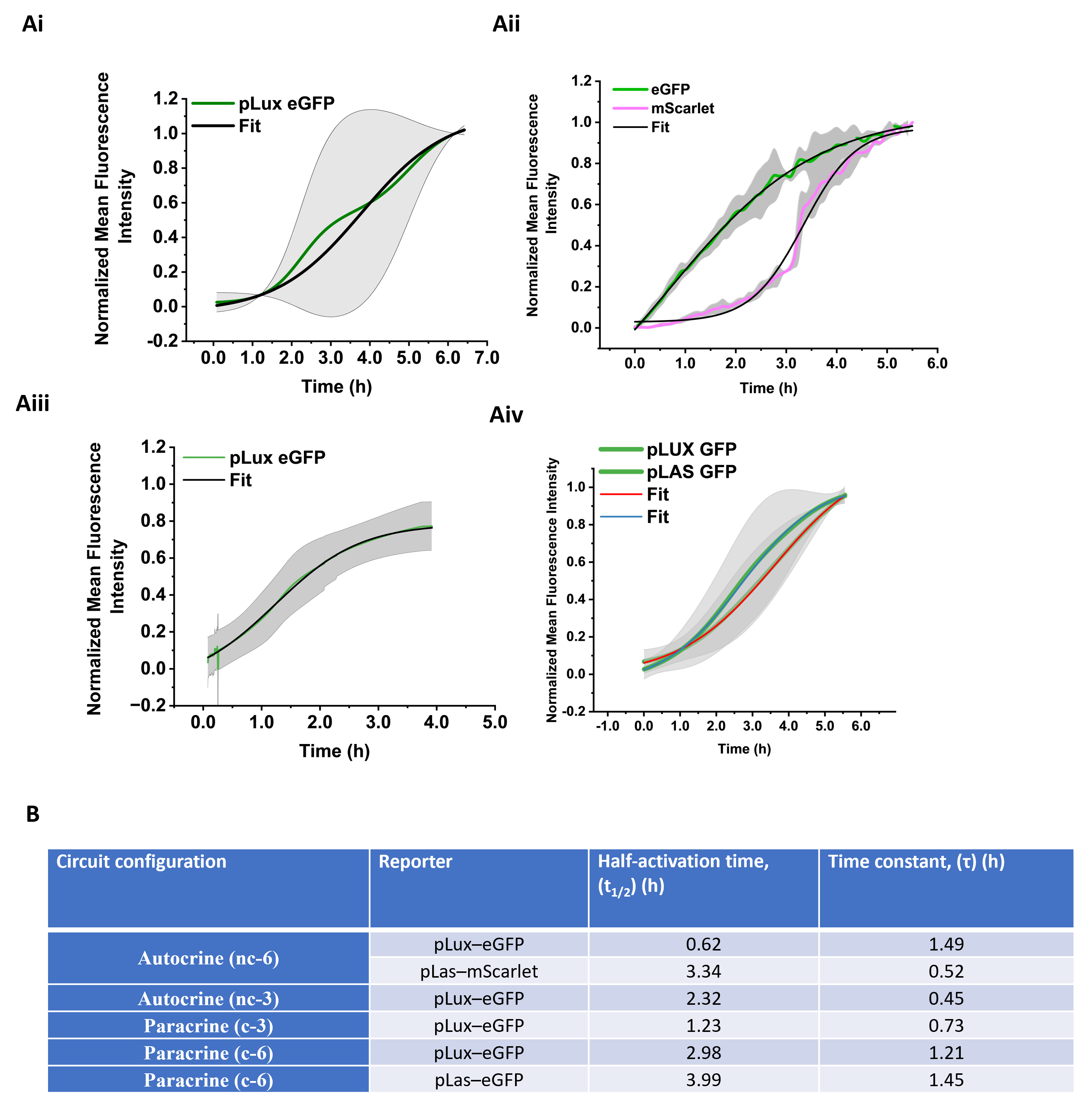

**Supplementary figure 8: Time evolution of gene expression in all four systems.**

Ai-Aiv: Four-parameter logistic (Boltzmann) fits (adjusted 𝑅^2^>0.99 R^2^>0.99) fo the gene expression curves for all the four systems. B: Table showing the summary of kinetic parameters extracted from four-parameter Boltzmann fits to the population-averaged fluorescence trajectories shown in Ai to Aiv.

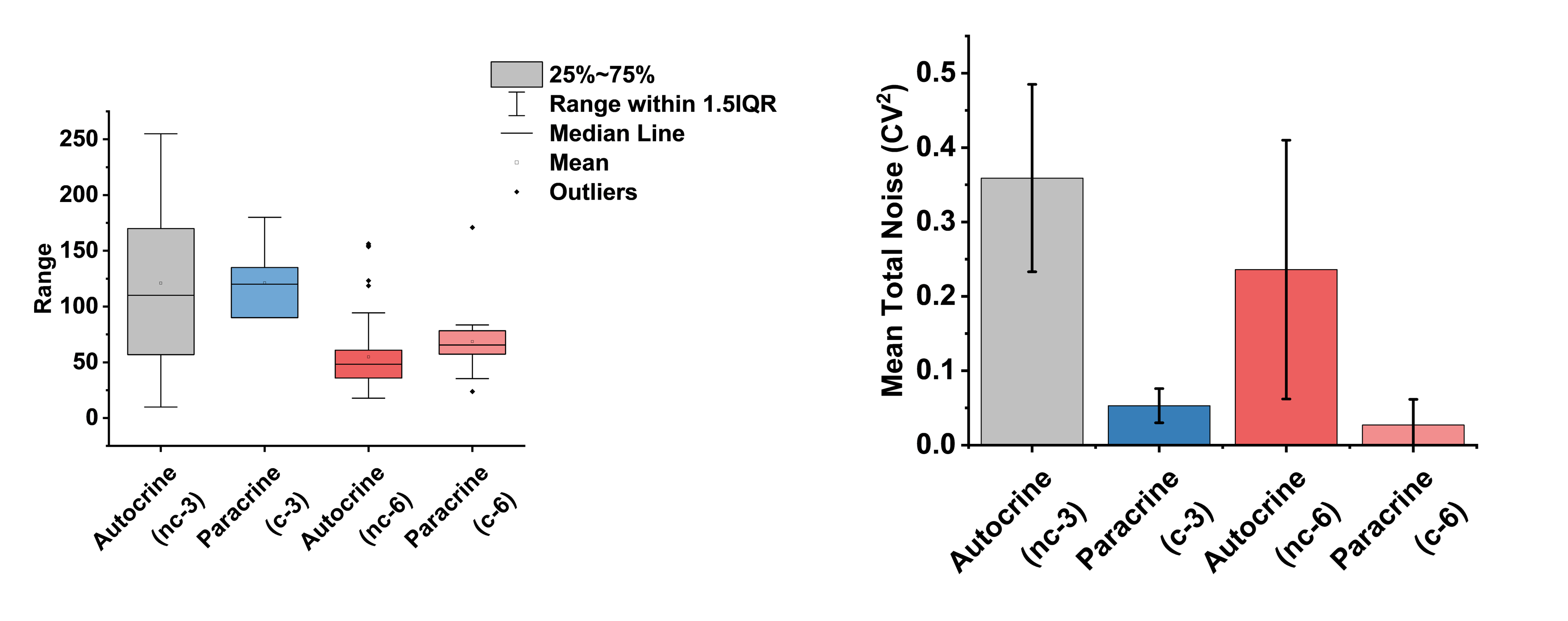

**Supplementary figure 9: Gene expression noise in all four systems.**

Left: Fluorescence intensity distribution of gene expression for all the four systems with two Autocrine and two Paracrine SCs populations. Boxplots at 4 nM plasmid concentration. Boxes indicate the 25–75% interquartile range, horizontal lines mark the median, open squares the mean, whiskers extend to 1.5× IQR, and symbols denote outliers. Right: Mean total noise (CV²) for the conditions in the left, preserves the hierarchy between architectures, with Paracrine systems remaining substantially less noisy than their Autocrine systems.

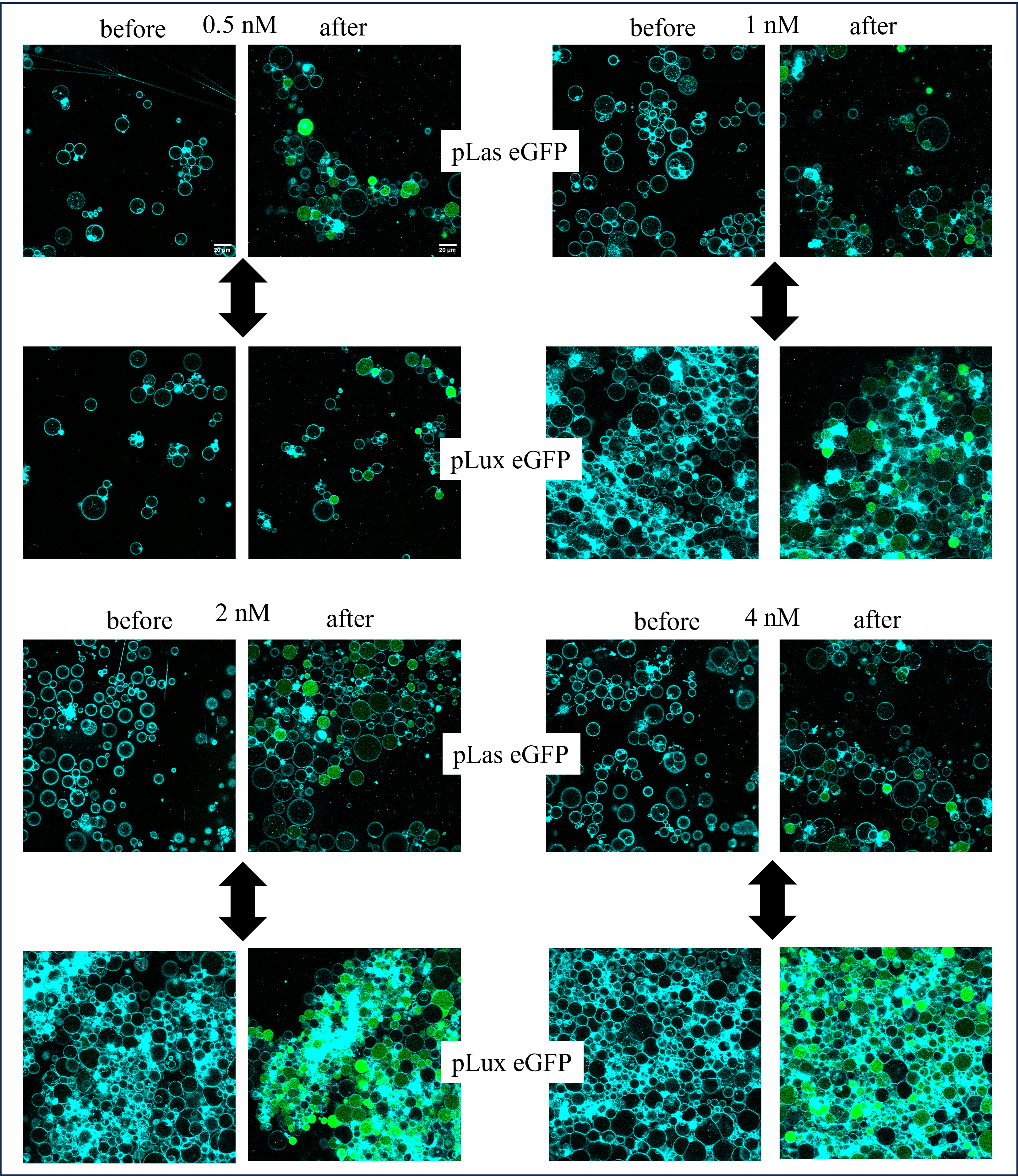

**Supplementary figure 10:** **Confocal microscopy images Paracrine (c-6) system with increasing plasmid concentration (0.5 nM to 4 nM).** Each concentration has pLas eGFP in the top panel expressing eGFP reporter protein and the bottom panel with pLux eGFP. Each panel also shows before and after images of the acquisition. The double headed arrow in the middle show, bidirectional nature of the communication. Insert in the top panel show scale bar, 20 µm.

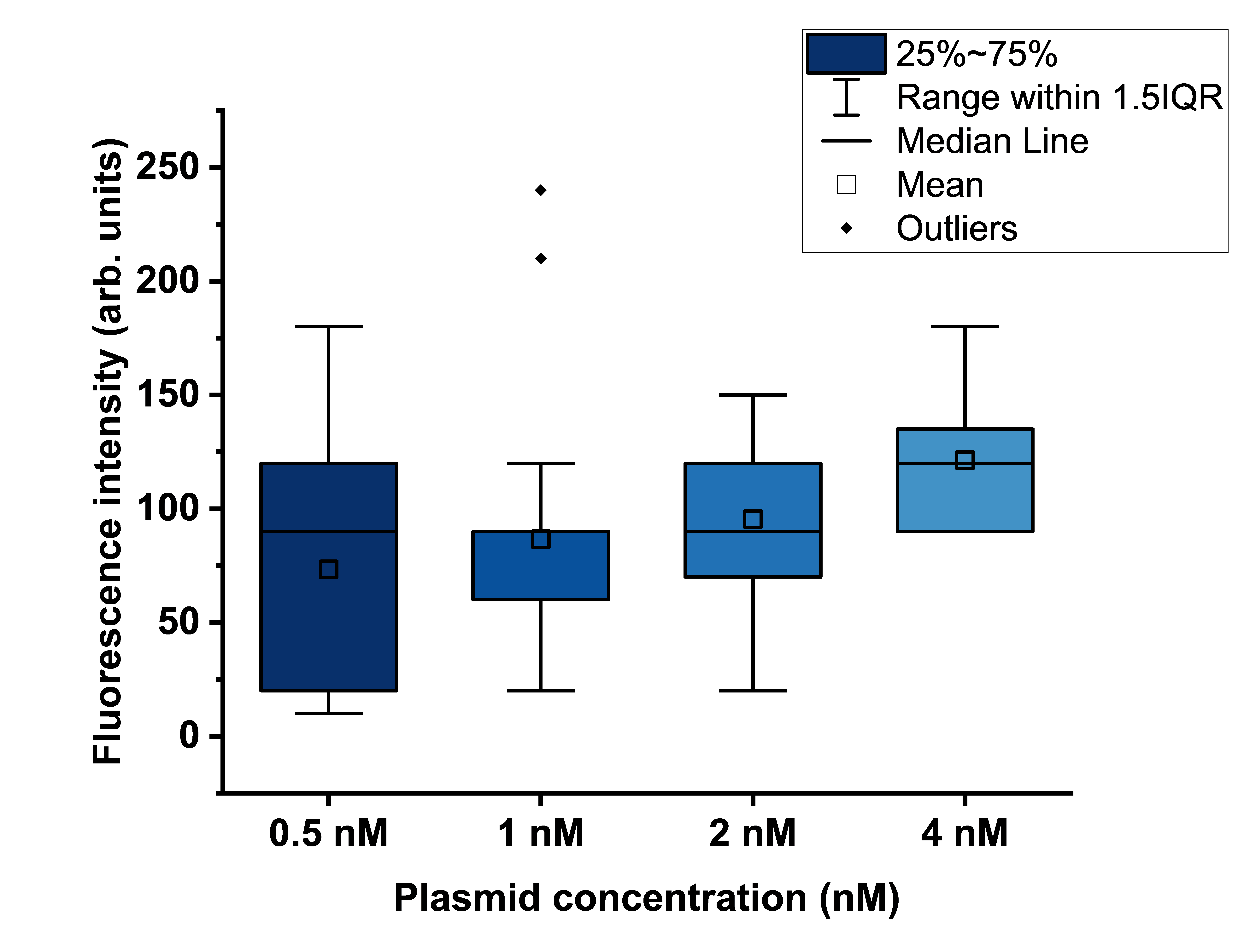

**Supplementary figure 11.** **Role of plasmid concentration in expression variability in uni-directional SCs populations.** Boxplots of single-vesicle pLux–eGFP reporter fluorescence in receiver synthetic cells of the uni-directional sender–receiver circuit at different plasmid concentrations (0.5, 1, 2, and 4 nM). Boxes indicate the 25–75% interquartile range, horizontal lines mark the median, open squares the mean, whiskers extend to 1.5× IQR, and symbols denote outliers.

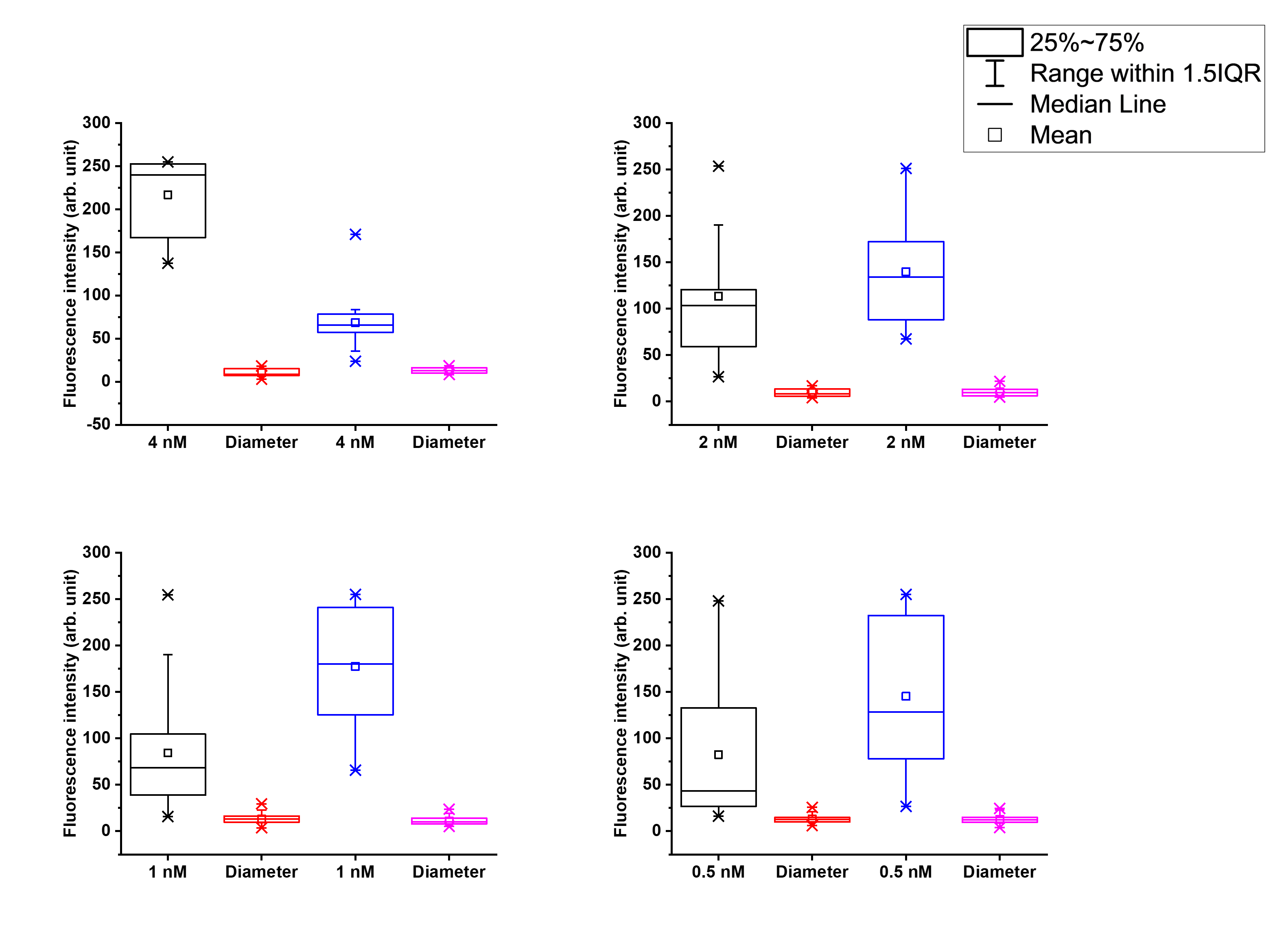

Supplementary figure 12. Role of plasmid concentration in expression variability in bi-directional SCs populations. Boxplots of single-vesicle endpoint measurements for the all-in-one cascade at different total plasmid concentrations. Each panel shows data for 4 nM (top left), 2 nM (top right), 1 nM (bottom left), and 0.5 nM (bottom right) plasmid DNA. Black boxplots report pLas–eGFP fluorescence intensities and blue boxplots report pLux–eGFP fluorescence intensities. Red and magenta boxplots show the corresponding vesicle diameters for the pLas and pLux populations, respectively. Boxes indicate the 25–75% interquartile range with medians (horizontal line), means (open squares), and whiskers extending to 1.5× IQR.

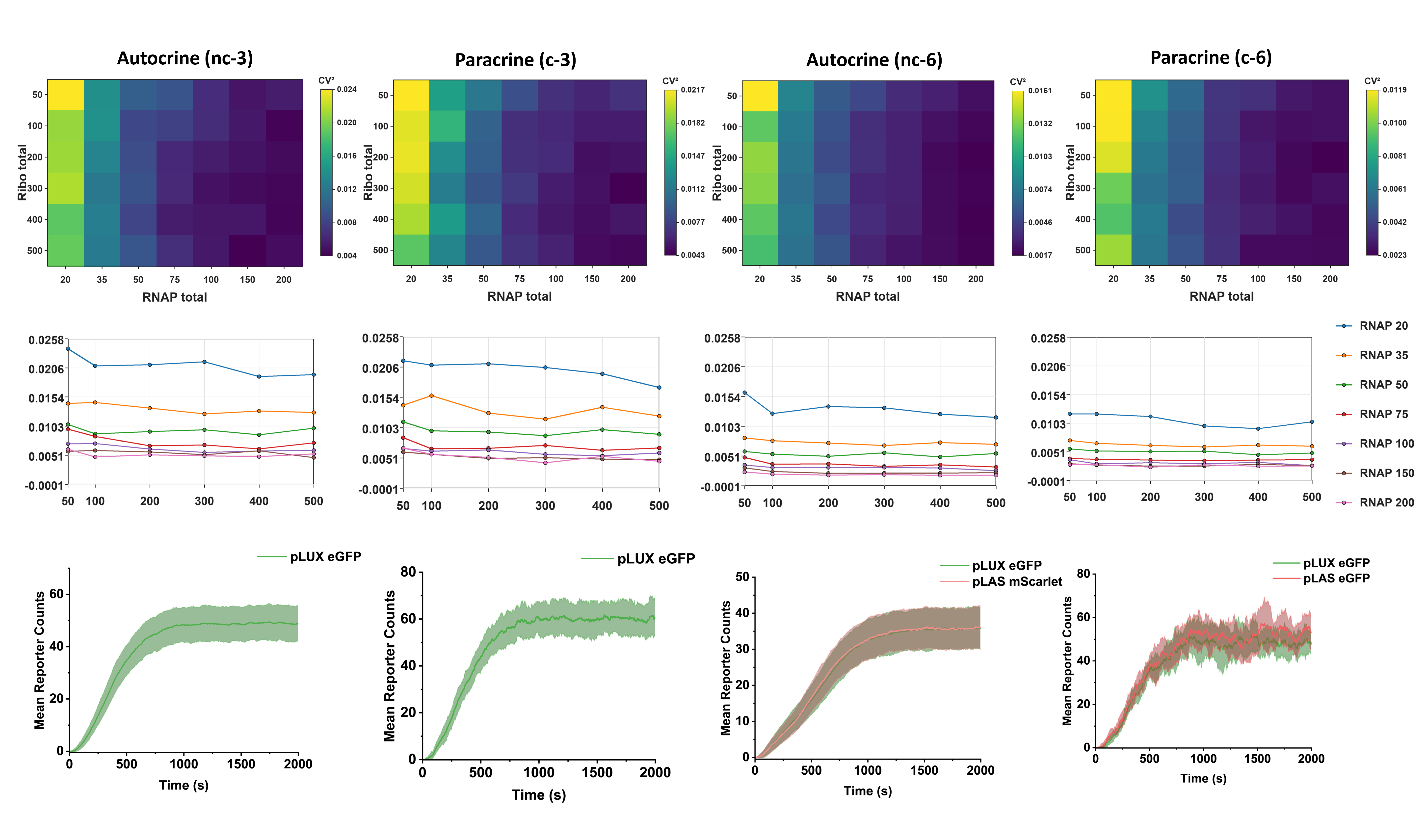

**Supplementary figure 13:** **Resource-dependent expression noise and kinetics in autocrine and paracrine synthetic-cell circuits.** Top row: Heat maps showing reporter noise, quantified as CV^2^, across total RNAP and ribosome availability. Middle row: Corresponding CV^2^ profiles as a function of ribosome total at different RNAP levels. Bottom row: Representative kinetic traces (N is 500) showing mean reporter counts over time for all four conditions. Autocrine three-component circuits showed reporter activation with resource-dependent noise, while paracrine receiver architecture generally maintained lower CV^2^ over a broad range of RNAP and ribosome levels. Shaded regions in kinetic traces represent variability around the mean.

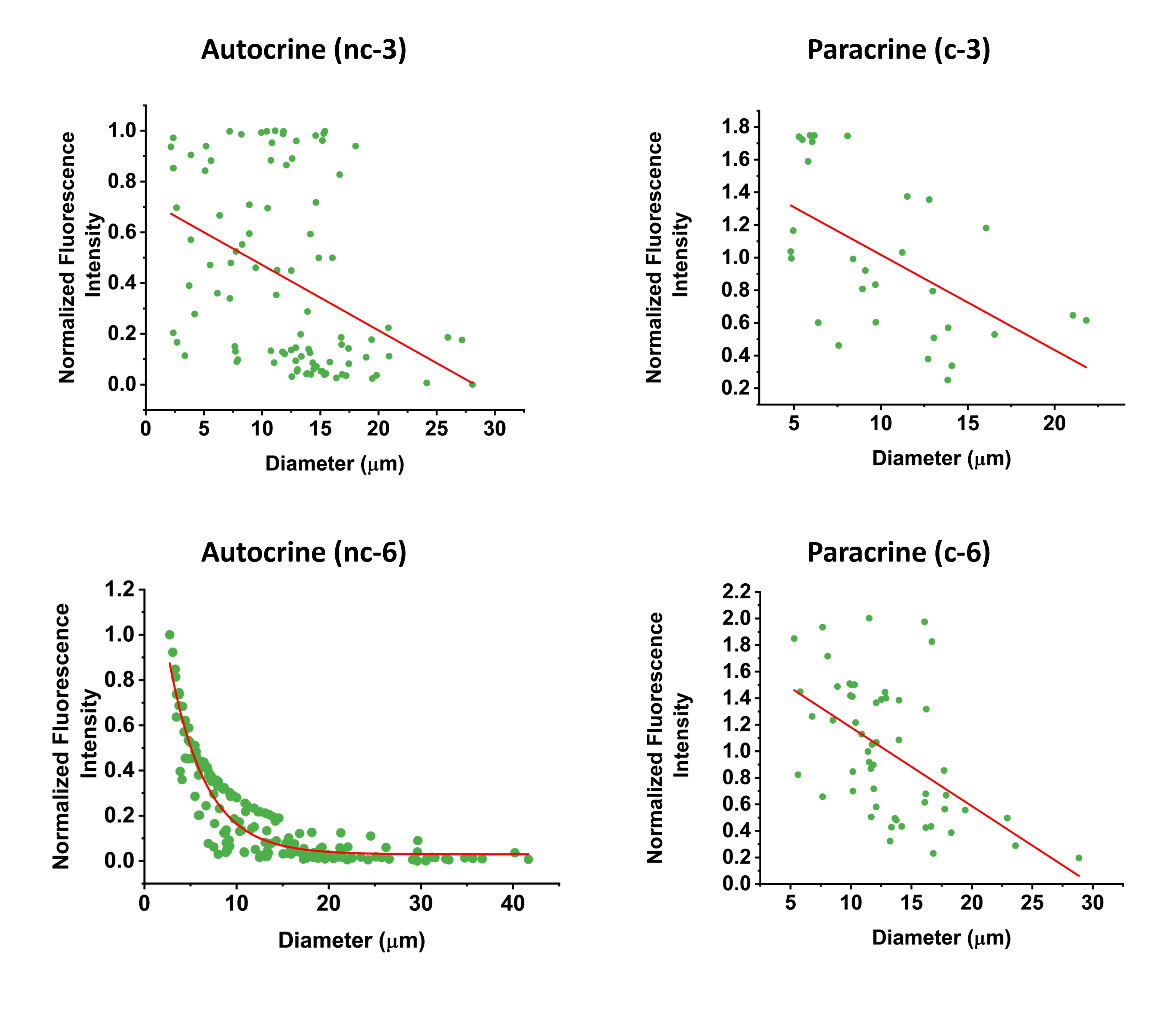

**Supplementary figure 14: Diameter-dependent reporter expression in autocrine and paracrine systems.** Scatter plots showing normalized fluorescence intensity as a function of synthetic-cell diameter for four circuit architectures: autocrine three-component circuit, paracrine three-component circuit, autocrine six-component circuit, and paracrine six-component circuit. Each green point represents an individual synthetic cell, and the red curve indicates the fitted size-dependence used for size correction (specified in the Methods). In all conditions, fluorescence intensity showed a negative dependence on vesicle diameter, indicating stronger reporter output in smaller compartments. The effect was most pronounced in the autocrine six-component circuit, where fluorescence decreased nonlinearly with increasing diameter. These fits were used to calculate size-adjusted residual fluorescence values and to estimate expression noise independently of vesicle size effects.

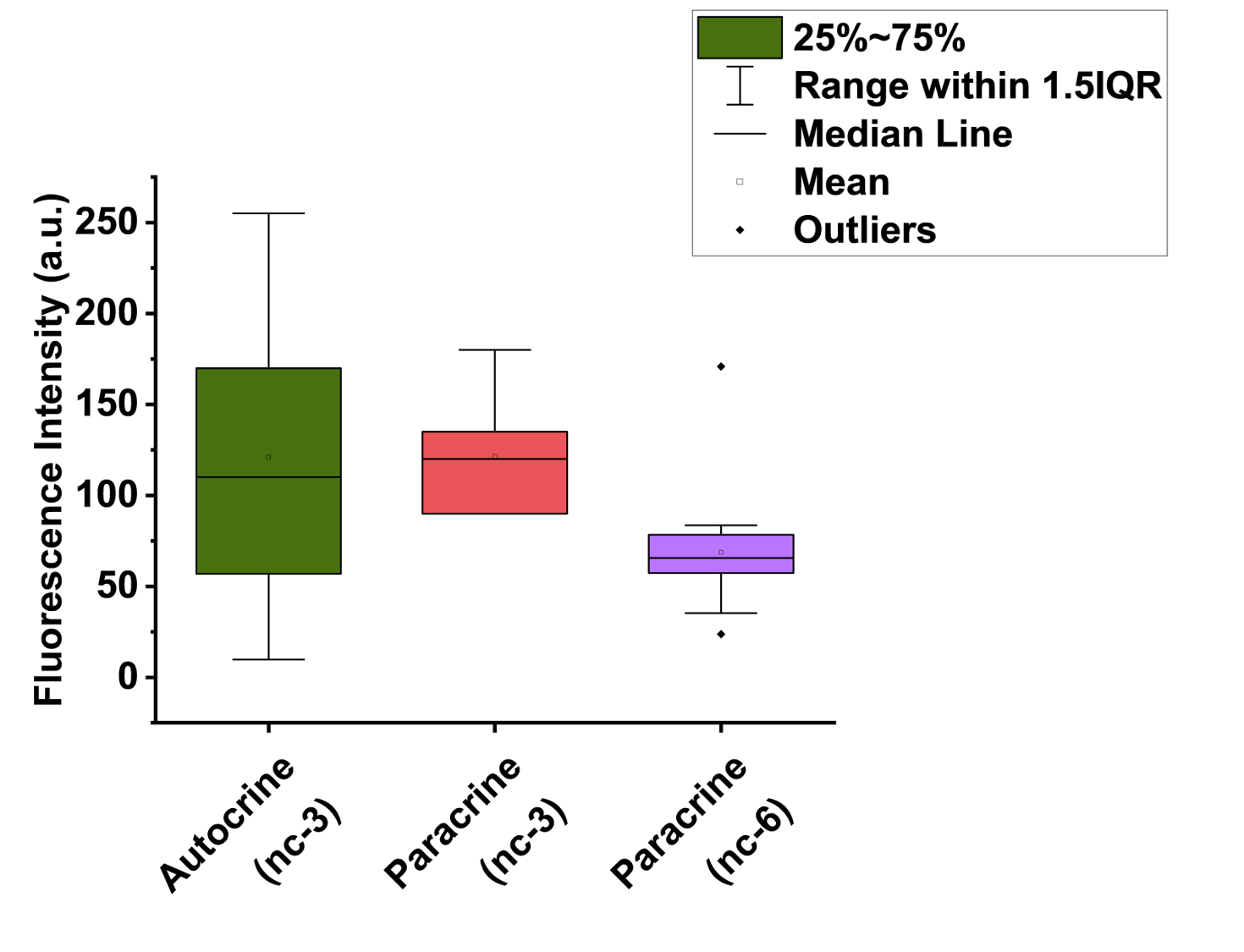

Supplementary figure 15. Comparative analysis of pLux–GFP variability under equi-resource conditions across SCs populations. Boxplots of single-vesicle pLux–eGFP fluorescence for non-communicating (green), uni-directional (red), and bi-directional (magenta) synthetic cells under equi-resource, 4nM plasmid concentration. Boxes show the 25–75% interquartile range, horizontal lines the median, open squares the mean, whiskers 1.5× IQR, and symbols outliers, highlighting the reduced fluorescence variability in communicating architectures.

Movie S1.

Timelapse video of eGFP and mScarlet expression in home-made CFES containing SCs. Scale bar – 20 µm

Movie S2.

Timelapse video of eGFP expression in home-made CFES containing SCs with non-communicating gene circuit. Scale bar – 20 µm
